# Genome diversification in globally distributed novel marine Proteobacteria is linked to environmental adaptation

**DOI:** 10.1101/814418

**Authors:** Zhichao Zhou, Patricia Q. Tran, Kristopher Kieft, Karthik Anantharaman

## Abstract

Proteobacteria constitute the most diverse and abundant group of microbes on Earth. In productive marine environments like deep-sea hydrothermal systems, Proteobacteria have been implicated in autotrophy coupled to sulfur, methane, and hydrogen oxidation, sulfate reduction, and denitrification. Beyond chemoautotrophy, little is known about the ecological significance of novel Proteobacteria that are globally distributed and active in hydrothermal systems. Here we apply multi-omics to characterize 51 metagenome-assembled genomes from three hydrothermal vent plumes in the Pacific and Atlantic Oceans that are affiliated with nine novel Proteobacteria lineages. Metabolic analyses revealed these organisms to contain a diverse functional repertoire including chemolithotrophic ability to utilize sulfur and C_1_ compounds, and chemoorganotrophic ability to utilize environment-derived fatty acids, aromatics, carbohydrates, and peptides. Comparative genomics with marine and terrestrial microbiomes suggests that lineage-associated functional traits could explain niche specificity. Our results shed light on the ecological functions and metabolic strategies of novel Proteobacteria in hydrothermal systems and beyond, and highlight the relationship between genome diversification and environmental adaptation.

## Introduction

Proteobacteria constitute one of the most diverse microbial phyla and are successful in most biomes on Earth. Proteobacteria are abundant from pole to pole in the world’s oceans^1, 2^, and also from the surface to the deep oceans in vertical cross-sections^3, 4^. Unlike other microbial lineages, Proteobacteria display an enormous functional repertoire and comprise phototrophs, autotrophs, and heterotrophs. In the surface oceans, heterotrophic proteobacteria such as SAR11, SAR86, and Roseobacter are abundant and highly successful bacterioplankton lineages which mainly rely on the availability and distribution of dissolved organic matter^5^. Phototrophic proteobacteria include groups of Alphaproteobacteria and Gammaproteobacteria that together comprise the group referred to as ‘marine aerobic anoxygenic phototrophs’ (AAnPs) that are capable of bacteriochlorophyll-dependent photosynthesis and can influence overall carbon and energy budgets in the oceans^6^. In the dark oceans, Proteobacteria drive carbon cycling through primary production associated with sulfur and methane oxidation^7^, as well as heterotrophy^8^. Given their abundance across marine environments and their wide range of metabolic traits, Proteobacteria represent an ideal lineage to investigate links between genome diversification and environmental adaptation. To constrain the environment in which to explore this question, we studied the distribution, metabolism, activity, and ecology of proteobacteria in deep-sea hydrothermal plumes, a system characterized by the presence of natural geochemical gradients.

Hydrothermal plumes are formed when hot fluids (up to 400°C) emanate from deep-sea hydrothermal vents and mix with cold deep ocean waters (2-4°C). This process causes steep thermal and chemical gradients at small spatial scales, and biotic and abiotic processes leading to the formation of a variety of ecological niches that can be exploited by microorganisms^9–11^. Hydrothermal fluids typically entrain substantial concentrations of reduced chemicals and substrates, e.g., hydrogen (H2), methane (CH4), hydrogen sulfide (H2S), ammonia (NH3), methanol, C1 compounds (formaldehyde, formate, and carbon monoxide), hydrocarbons and metals (Fe, Mn, As, and etc) ^12,13–20^. Hydrothermal biogeochemistry strongly influences microbial community composition in the plume and surrounding environments^10^. In chemosynthetic environments of the oceans such as hydrothermal systems, proteobacterial groups actively participate in primary production by utilizing a wide variety of reduced substrates^10, 21^. Specific examples include diverse and active populations of *Sulfurimonas* and *Sulfurovum* (Epsilonproteobacteria) species that oxidize reduced sulfur compounds; *Thioglobus/*SUP05 and *Beggiatoa* (Gammaproteobacteria) that oxidize reduced sulfur compounds and hydrogen for energy generation^10^; and *Methylophaga* and Methylococcaceae (Gammaproteobacteria) that can oxidize methane, methanol, and hydrocarbons^22^.

Besides plumes, hydrothermal systems host several chemosynthetic environments dominated by Proteobacteria. Thermophilic Epsilonproteobacteria belonging to the genera *Nautilia* and *Lebetimonas* that can oxidize hydrogen with elemental sulfur, are prevalent on the hydrothermal vent deposits^23^. On the periphery of the venting area with lower temperatures (< 55°C), vent animals (such as tubeworms, crabs, bivalves) depend on the chemosynthetic symbiotic Proteobacteria as the source of organic carbon and energy^10^. Typically, proteobacterial endosymbionts (mostly Gammaproteobacteria) of tubeworms can oxidize reduced sulfur species^24^, while proteobacterial endosymbionts of bivalves can perform oxidation of reduced sulfur, methane, hydrogen and carbon monoxide^24–26^. Beyond these host animals, little is known about whether other microbes could also utilize organic compounds from vent-derived chemosynthesis^9^. Organic carbon from primary production may be used in heterotrophy in hydrothermal plumes as they disperse or even be consumed locally in vent-associated environments. However, due to the predisposition of hydrothermal plumes as chemosynthetic environments, microorganisms especially bacteria associated with heterotrophy in plumes remain little-studied. Organisms in deep-sea systems are often versatile and can exhibit mixotrophic characteristics. To elucidate the versatile metabolism, evolutionary relationships, and ecology of abundant groups of little-characterized deep-sea Proteobacteria, we reconstructed and compared genomes, transcriptomes, and protein families to Proteobacteria from other terrestrial and marine biomes.

In this study, we reconstructed 51 novel proteobacterial genomes from the deep-sea hydrothermal plumes and surrounding background seawaters at three distinct locations, namely Guaymas Basin and Lau Basin in the Pacific Ocean, and Mid-Cayman Rise in the Atlantic Ocean. These hydrothermal systems comprise gradients in ocean depth (∼1900m to 5000m), geophysical parameters (back-arc basin to sedimented hydrothermal system), and chemistry^27–29^. Our genome-resolved metagenomics approach has allowed the discovery of novel microbial groups of Proteobacteria based on phylogeny reconstructed using both concatenated ribosomal proteins and 16S rRNA genes and identified metabolic characteristics of specific proteobacterial organisms and lineages. Metatranscriptomics-derived measurements enabled us to describe and compare the activity of novel Proteobacteria across a range of environments within and between different plumes and deep ocean samples. The ecological and functional knowledge associated with the metabolism of these novel microbial groups provides insights into organic carbon metabolism, energy transformations, and adaptive strategies in hydrothermal vent ecosystems and beyond. While primarily found in hydrothermal systems, these novel Proteobacteria have a worldwide distribution and can be observed outside of marine environments including freshwaters and the terrestrial subsurface. Overall, our study reveals that genome diversification in globally prevalent and abundant Proteobacteria is associated with environmental adaptation and suggests that the distribution of functional traits could potentially explain their niche-adapting mechanisms.

## Results

### Sampling and reconstruction of genomes from deep-sea hydrothermal plumes

Hydrothermal vent plume and background deep-sea samples were acquired during the following cruises: R/V New Horizon to Guaymas Basin (July 2004), R/V Atlantis to Mid-Cayman Rise (Jan 2012 and Jun 2013) for Cayman Deep (*Piccard*) and Shallow (*Von Damm*) plume and background seawater samples, and R/V Thomas G Thompson to the Lau Basin (May-Jul 2009) for both plume and background seawater samples. Details of sample collection, preservation, and DNA/RNA extraction and processing are described in detail elsewhere^9, 27, 29^. Briefly, samples were collected using shipboard filtration (Guaymas) or filtered *in situ* using the SUPR sampler mounted on a remotely operated vehicle^27, 29^.

In this study, we reconstructed genomes from publicly available shotgun metagenomic sequencing datasets from 19 deep-sea hydrothermal plume and surrounding background seawater samples from Guaymas Basin (Guaymas), Mid-Cayman Rise (Cayman) and Lau Basin (Lau) (Fig. 1). Additionally, we analyzed the 14 metatranscriptomic sequencing datasets that were paired with metagenomics samples from Guaymas and Cayman (Supplementary Table S1). Following quality-control, filtered reads were used to assemble scaffolds *de novo* according to the location of metagenomic samples, which includes one vent from Guaymas, two vents from Cayman (*Piccard* and *Von Damm*) and five vents from Lau (Abe, Kilo Moana, Mariner, Tahi Moana, and Tui Malia). Application of the metagenomic binning approach resulted in 250 metagenome-assembled genomes (MAGs) which have genome completeness > 50% and genome contamination/redundancy < 10% and were used for downstream analyses.

**Figure 1.**
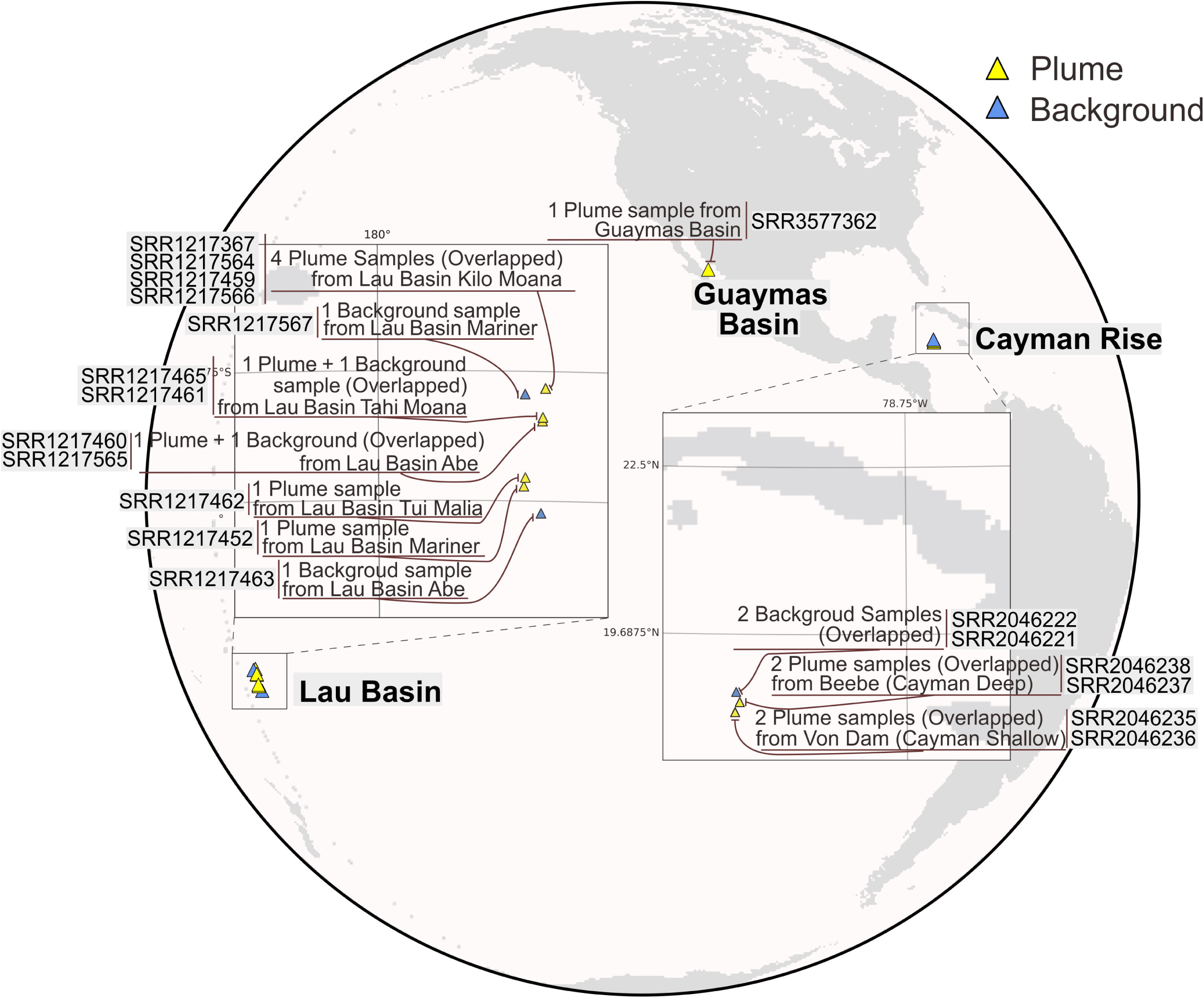
Schematic map representing the sampling locations of hydrothermal samples. The retrieved metagenomic datasets include one from Guaymas Basin, twelve from Lau Basin, and six from Mid-Cayman Rise. Detailed sample, metagenome, and metatranscriptome information is labeled.

### Phylogeny and identification of novel proteobacteria groups

To identify the taxonomy of the reconstructed genomes, we used a comprehensive phylogenetic approach. First, we constructed a phylogenetic tree using a set of concatenated 16 ribosomal proteins (RP16 tree). The reconstructed phylogenetic tree revealed that bacteria comprised 219 of the 250 genomes, with 92 of them from the group Proteobacteria. We then conducted detailed taxonomic curation of all reconstructed genomes by comparison with specific databases and phylogenetic trees, namely NCBI, GTDB^30^, and RP16. A companion 16S rRNA gene phylogenetic tree (using genes retrieved from MAGs) also has the congruent phylogeny (Supplementary Fig. S1). Of the 92 Proteobacteria genomes, we determined 51 to be phylogenetically novel (Supplementary Table S2); all lack a defined taxonomy at the scale of family, order, and/or phylum, coupled with a lack of understanding of their metabolism and ecology.

Based on the RP16 tree, we classified and defined 9 proteobacterial groups at different levels, including 2 phyla, 3 classes and 4 families (Fig. 2 and Supplementary Fig. S2, S3). We propose the names Marenostrumaceae for UBA2165 (since the UBA2165 type strain was first reconstructed from the Mediterranean Sea^31^; known as ‘Mare Nostrum’ in Latin); Hyrcanianaceae for group Casp-alpha2 (since the Casp-alpha2 type strain was first reconstructed from the Caspian Sea^31^; known as the ‘Hyrcanian Ocean’ in ancient Greek), Taraoceanobacteraceae for UBA11654, which was first reconstructed from Tara Ocean metagenome datasets^32^; Riflewellaceae for UBA4486 which was first described from terrestrial aquifer wells at Rifle, Colorado, USA^33^; Marinioligotrophales for the formerly OMG bacteria, the Oligotrophic Marine Gammaproteobacteria^34^; Planktothermales for UBA7887, which are reconstructed from hydrothermal plume environments^31^; Thalassomicrobiales for the SAR86 bacteria, which are ubiquitously distributed in oceans^35^; and Kappaproteobacteria for the former LS-SOB group^31^, which are ubiquitous in coastal systems and the ocean water column. This classification was also supported by the taxonomic tree associated with the GTDB^30^ taxonomy database (https://gtdb.ecogenomic.org/tree). Finally, Lambdaproteobacteria was first discovered and characterized from a groundwater ecosystem^36^, while this is the first study to retrieve and study genomes recovered from deep-sea hydrothermal plumes or any marine environment for this novel phylum. The general functional trait profile on investigating the most comment genetic property indicates that most of the novel proteobacterial genomes from this study have metabolic capacities associated with aerobic respiration, sulfur cycling, and CO2 fixation (Fig. 2 and Supplementary Table S3).

**Figure 2.**
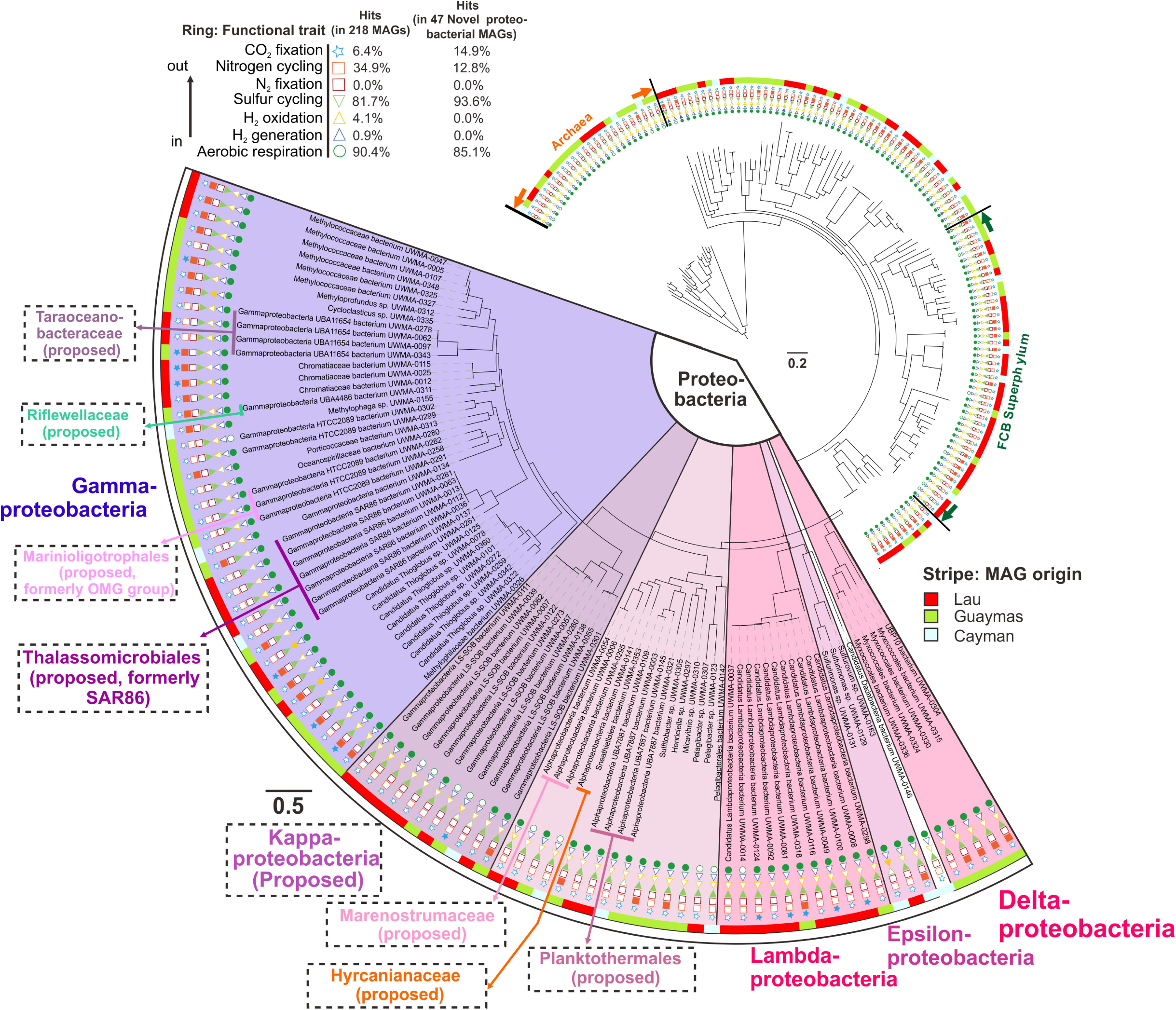
Phylogenetic tree of the hydrothermal plume and deep-sea-derived MAGs based on the concatenated 16 ribosomal protein alignment (RP16 tree. Functional traits associated with carbon, nitrogen, sulfur, hydrogen cycling, and oxygen respiration are shown. Filled and unfilled circles denote the presence/absence of function traits within a genome. This tree was visualized using iTOL^89^ (https://itol.embl.de/).

### Distribution and abundance

To understand the biogeography and importance of Proteobacteria in hydrothermal environments and beyond, we estimated their distribution and abundance. Novel proteobacterial genomes identified by us constituted 23% of all reconstructed genomes from the hydrothermal environments studied here. Yet, analyses of genome coverage indicated that they comprised 36% of the microbial community, which suggests they are more abundant than other microbial groups (Supplementary Table S4). To determine if this abundance of novel Proteobacteria also translated to a higher activity of these organisms in hydrothermal plumes, we analyzed gene expression using metatranscriptomics. Similar to observations from metagenomics, metatranscriptomic data suggests that these novel Proteobacteria had higher activity in comparison to other microbial groups. Specifically from Guaymas Basin, novel Proteobacteria constituted 17% of the microbial community by abundance yet accounted for 20% of all microbial activity. Additionally, the proportion of gene expression (cDNA) among all novel Proteobacteria genomes was higher than abundant genomes representing organisms from dominant marine phyla such as Chloroflexi (SAR202) and Bacteroidetes (Supplementary Table S4).

We then assessed the global distribution of novel Proteobacteria by examining their presence in publicly available data in the Integrated Microbial Genomes and Metagenomes database (IMG/M)^37^. While most novel proteobacterial groups were widely distributed, specifically Kappaproteobacteria, Thalassomicrobiales, and Marinioligotrophales were especially abundant and distributed worldwide in oceanic and coastal environments (Fig. 3). In addition, these groups were also observed in other environments outside of marine systems such as in association with a host (symbiosis), terrestrial environments, and engineered systems indicating their widespread distribution in diverse eco-niches (Fig. 3).

**Figure 3.**
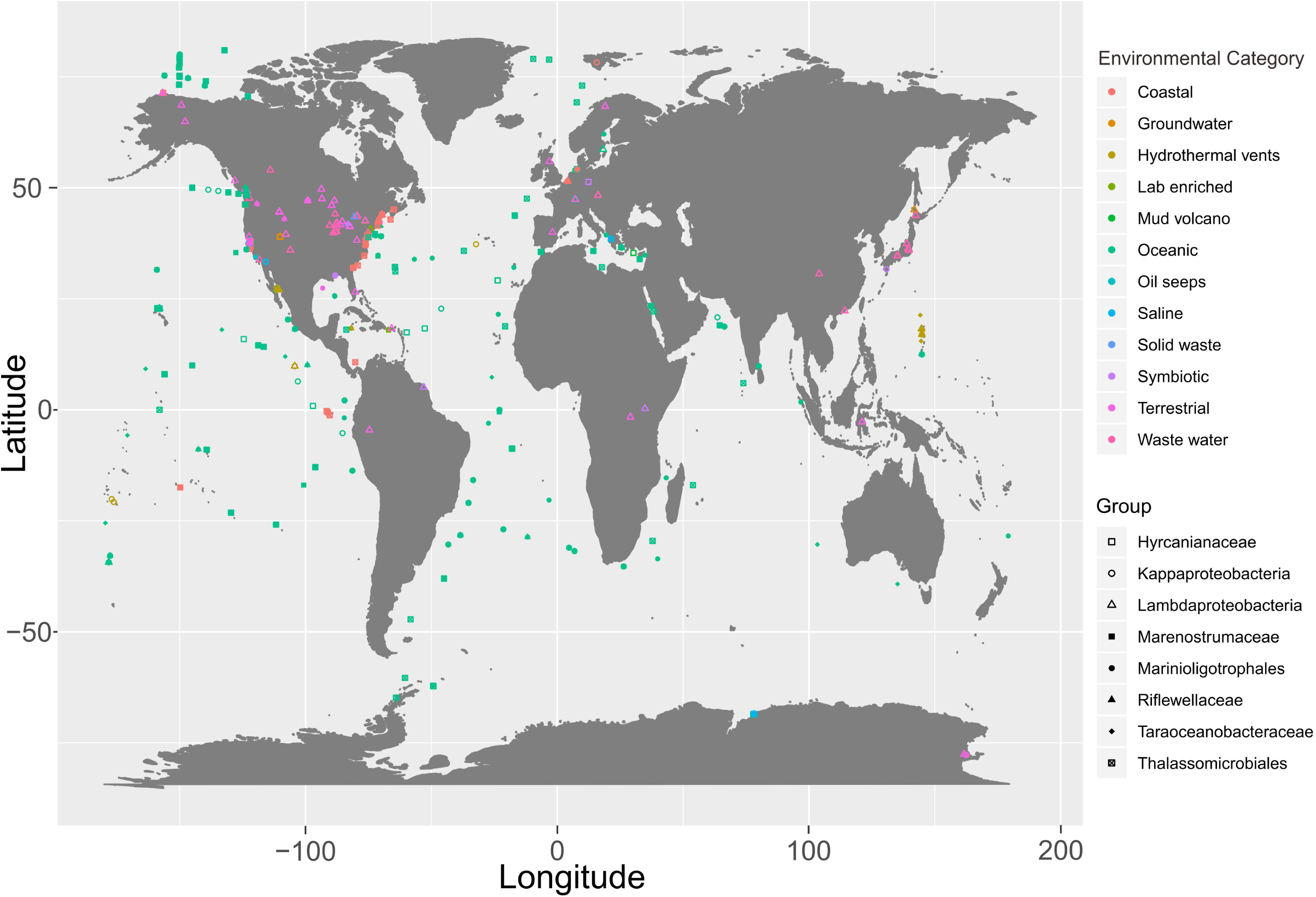
World map showing the distribution of nine novel proteobacterial lineages. Environmental categories were parsed and summarized from “ecosystem type” series information from all investigated IMG metagenomes.

### Central metabolism and respiration

We chose genomes from this study and publicly available genomes with completeness over 80% to reconstruct the genome functional profiles, in order to reflect the general metabolic and functional capacities of these novel proteobacterial groups (Supplementary Tables S5, S6). Additionally, the metatranscriptomic datasets from Guaymas and Cayman enabled investigation of gene expression associated with metabolic pathways involved in element and energy cycling (Figs. 4, 5 and Supplementary Table S6).

**Figure 4.**
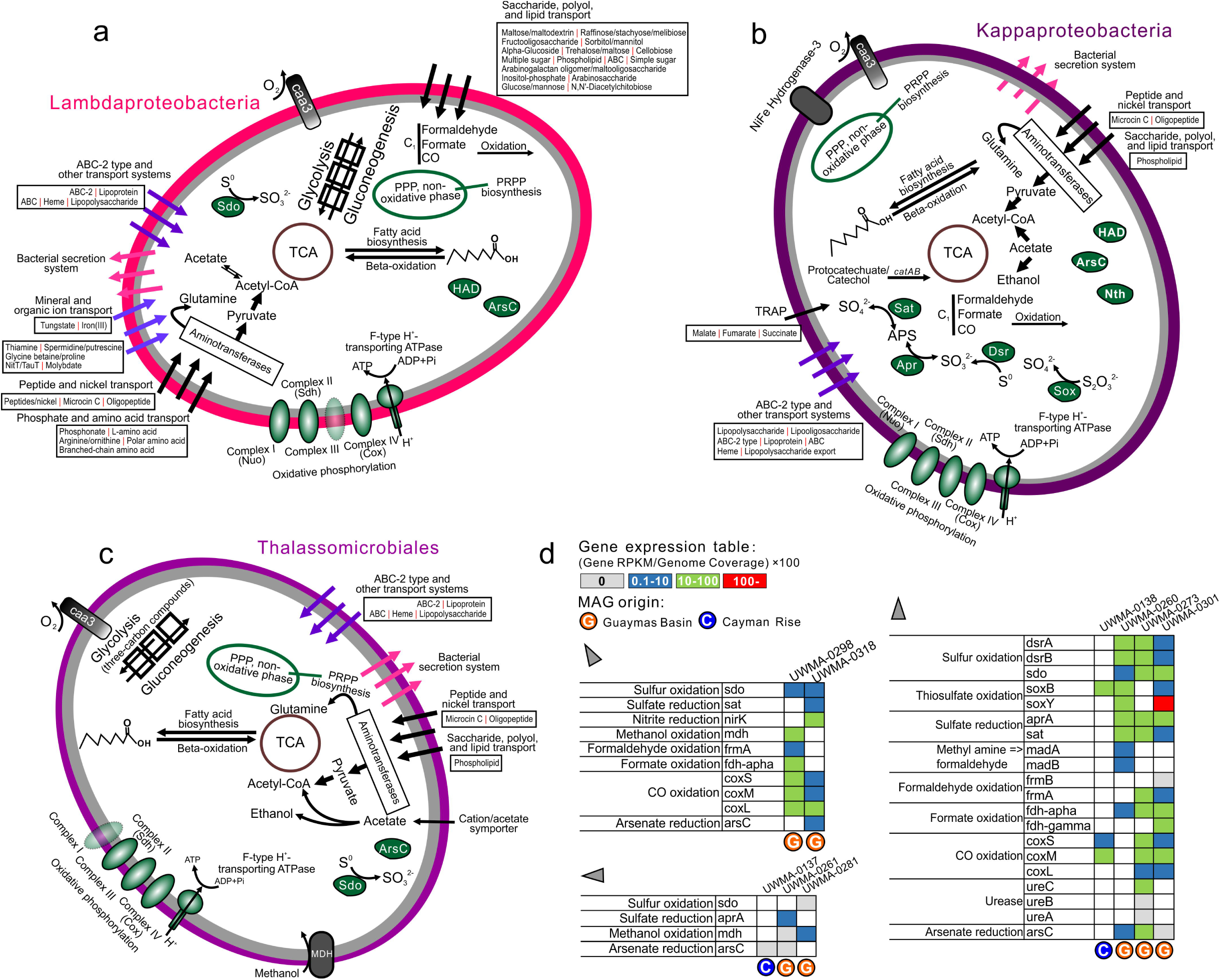
Cellular maps of inferred metabolic capacities and activities of Lambdaproteobacteria, Thalassomicrobiales, and Kappaproteobacteria. **a-c.** Organismal profile of metabolic capacity for Lambdaproteobacteria, Kappaproteobacteria, and Thalassomicrobiales, respectively. Items in grey indicate traits that are not present in over 50% of sampled genomes. **d.** Tables indicating the normalized gene expression level associated with important pathways associated with energy metabolism in novel Proteobacteria genomes.

**Figure 5.**
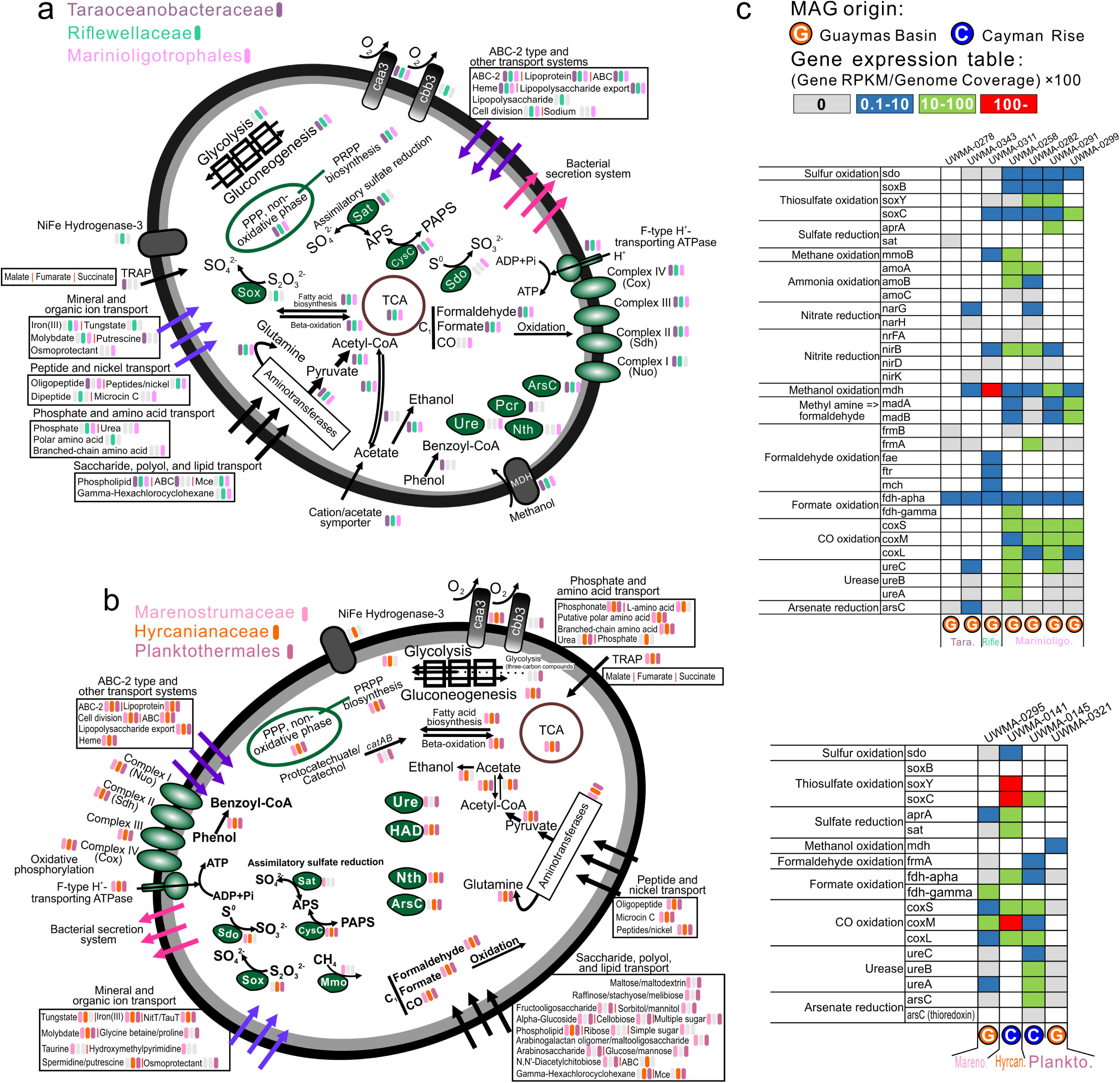
Cellular maps of inferred metabolic capacities and activities of the six novel proteobacterial families and orders. **a.** Organismal profiles of metabolic capacities for Taraoceanobacteraceae, Riflewellaceae, Marinioligotrophales, **b.** Organismal profiles of metabolic capacities for Marenostrumaceae, Hyrcaniceae, Planktothermales. Items in grey indicate traits that are not present in over 50% of sampled genomes. **c.** Tables indicating the normalized gene expression level associated with important pathways associated with energy metabolism in novel Proteobacteria genomes.

All proteobacterial genomes contained genes for central carbon metabolism pathways for biosynthesis and energy transfer: citric acid cycle (TCA cycle), glycolysis (some could only metabolize three-carbon compounds) and gluconeogenesis, peptide and amino acid utilization, pentose phosphate pathway (PPP pathway) and PRPP biosynthesis for the generation of some nucleotide and amino acid precursors, and fatty acid biosynthesis and beta-oxidation (Fig. 4, 5 and Supplementary Table S7). All novel proteobacterial groups contain either caa3/cbb3 type cytochrome *c* oxidases for aerobic respiration and nearly a complete set of complexes for oxidative phosphorylation for the generation of proton motive force dependent-ATP synthesis, suggesting that fermentation and respiration can both take place.

### Organic carbon metabolism

We investigated organic carbon metabolism in novel proteobacteria by identifying the presence of transporters, secretion systems, carbohydrate-active enzymes (CAZYmes)^38^, and peptidases in the genomes. Functional predictions of cell membrane transport and secretion systems indicate that Lambdaproteobacteria, Marenostrumaceae, and Planktothermales have more transporters involved with monosaccharides, polysaccharides, polyols, and lipids which may be used as organic nutrients for carbon and energy cycling. Moreover, Kappaproteobacteria, Taraoceanobacteraceae, Marenostrumaceae, Hyrcanianaceae, and Planktothermales contain tripartite ATP-independent periplasmic transporters for transporting C4-dicarboxylates^39^ while Thalassomicrobiales, Taraoceanobacteraceae, Riflewellaceae, and Marinioligotrophales contain cation/acetate symporters for direct acetate incorporation (Figs. 4, 5). Our analyses indicate that many of these organisms are capable of fermentation from acetate to ethanol and have wide-ranging capacities for the oxidation of C1 compounds such as formaldehyde, formate, and carbon monoxide. Meanwhile, some of them can also degrade and utilize aromatic compounds, such as phenol and protocatechuate/catechol (Figs. 4, 5). Except for Thalassomicrobiales, Taraoceanobacteraceae, and Riflewellaceae, all other groups contain organisms that can actively oxidize C1 compounds from hydrothermal environments. All groups except for Kappaproteobacteria, Marenostrumaceae, and Hyrcanianaceae demonstrated high levels of activity associated with methanol oxidation. The highest activity was observed in Riflewellaceae organisms (Figs. 4, 5). Genomes from Kappaproteobacteria and Marinioligotrophales encoded highly active genes for the utilization of methyl amines. These results indicate that these Proteobacteria are well adapted to hydrothermal ecosystems, and their metabolic activities are connected to the transformation of organic compounds of hydrothermal origin which are entrained in the plume and surrounding environments, such as C1 (formate, formaldehyde, carbon monoxide)^13–16^ and methylated compounds (methanol and methylamine)^17, 18^.

To study the potential of novel proteobacteria to breakdown carbohydrates and proteins, we screened all genomes for the presence of CAZymes and peptidases. Patterns of normalized CAZyme and peptidase gene coverage demonstrate that Lambdaproteobacteria, Kappaproteobacteria, and Marinioligotrophales are involved in carbohydrate and protein scavenging (Figs. 6, 7). Gene expression (cDNA) profiles suggest that Lambdaproteobacteria, Kappaproteobacteria, and Marinioligotrophales CAZymes and peptidases are highly active in Guaymas and Cayman (Supplementary Tables S9, S10). GH109 (α-N-acetylgalactosaminidase) for degradation of amino sugars, GH23 (lysozyme) for degradation of peptidoglycans and PL22 (oligogalacturonide lyase) for degradation of oligogalacturonides (a product of pectin degradation) are widely distributed in the Proteobacteria. In contrast, other families have limited distributions in specific lineages. For example, GH13 (α-amylase) for degradation of starch and pullulans, GH38 (α-mannosidase) and GH76 (ß-mannosidase) for degradation of mannooligosaccharides, and GH74 (ß-1,4-endoglucanase, endoglucanase) for degradation of cellulose and xyloglucan are primarily distributed in Lamdaproteobacteria and highly expressed. The other five novel proteobacterial groups (Thalassomicrobiales, Taraoceanobacteraceae, Riflewellaceae, Hyrcanianaceae, and Planktothermales) exhibited a limited distribution of CAZymes and peptidases suggesting little to no involvement in carbohydrate and protein breakdown. Overall, this indicates that novel proteobacteria participate in carbon and energy cycling in hydrothermal environments with differing metabolic strategies, which are mainly reflected in their divergent organotrophic capacities.

Lambdaproteobacteria genomes encoded the highest abundance of peptidases (Figs. 6, 7). The most abundant and actively expressed peptidases were related to protein quality control and regulation, e.g., S16 (Lon-A peptidase), an unfolded protein degrader, M41 (FtsH peptidase) and I87 (FtsH inhibitor), which inhibit FtsH and modulate the degradation of mistranslation products that might disrupt membranes, and S14 (Clp peptidase), which regulates specific protein degradation (Fig. 6 and Supplementary Table S9). This could result from stress response activities that may be induced from association with high-temperature fluids^40^ and serve as a protective and regulatory mechanism for cell membrane maintenance and protein transformations. Besides, there are also other abundant and active endo/extracellular peptidases in Lambdaproteobacteria which are responsible for harvesting and degrading peptides and endopeptide turnover, e.g., C26 (gamma-glutamyl hydrolase) for the turnover of folyl poly-gamma-glutamates, S33 (prolyl aminopeptidase), an extracellular peptidase prone to proline-rich substrate utilization, M20A and M20D (glutamate carboxypeptidase), and M23B (ß-lytic metallopeptidase) which lyse cell walls.

**Figure 6.**
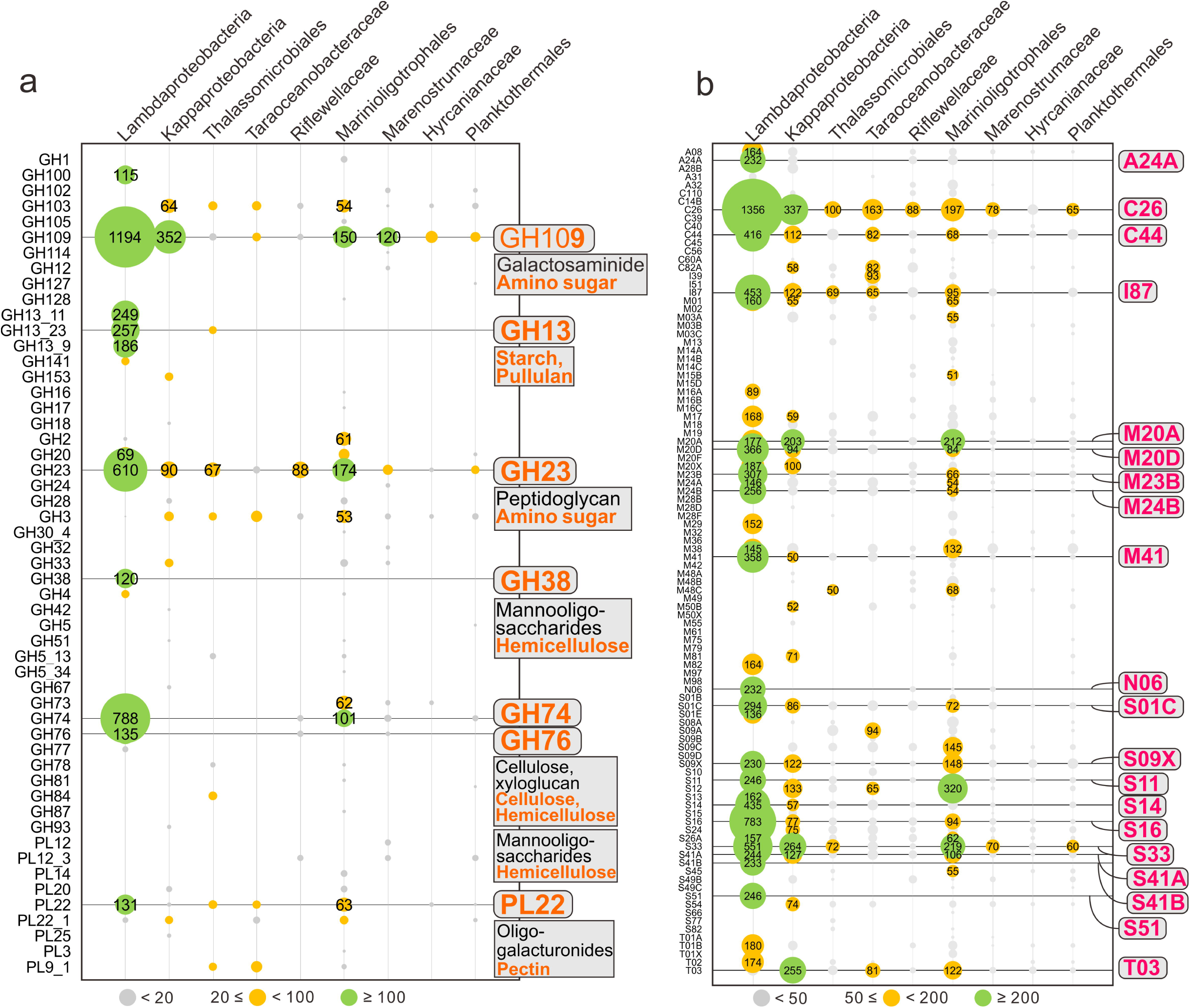
Coverage profiles of carbohydrate-active enzymes (CAZymes) and peptidases in novel proteobacterial genomes. **a.** Glycoside Hydrolase (GH) and Polysaccharide Lyase (PL) gene coverage was calculated by multiplying the identified number of CAZymes with normalized genome coverage. **b.** Peptidase (also includes peptidase inhibitors) gene coverage was calculated by multiplying the identified number of peptidases with normalized genome coverage.

**Figure 7.**
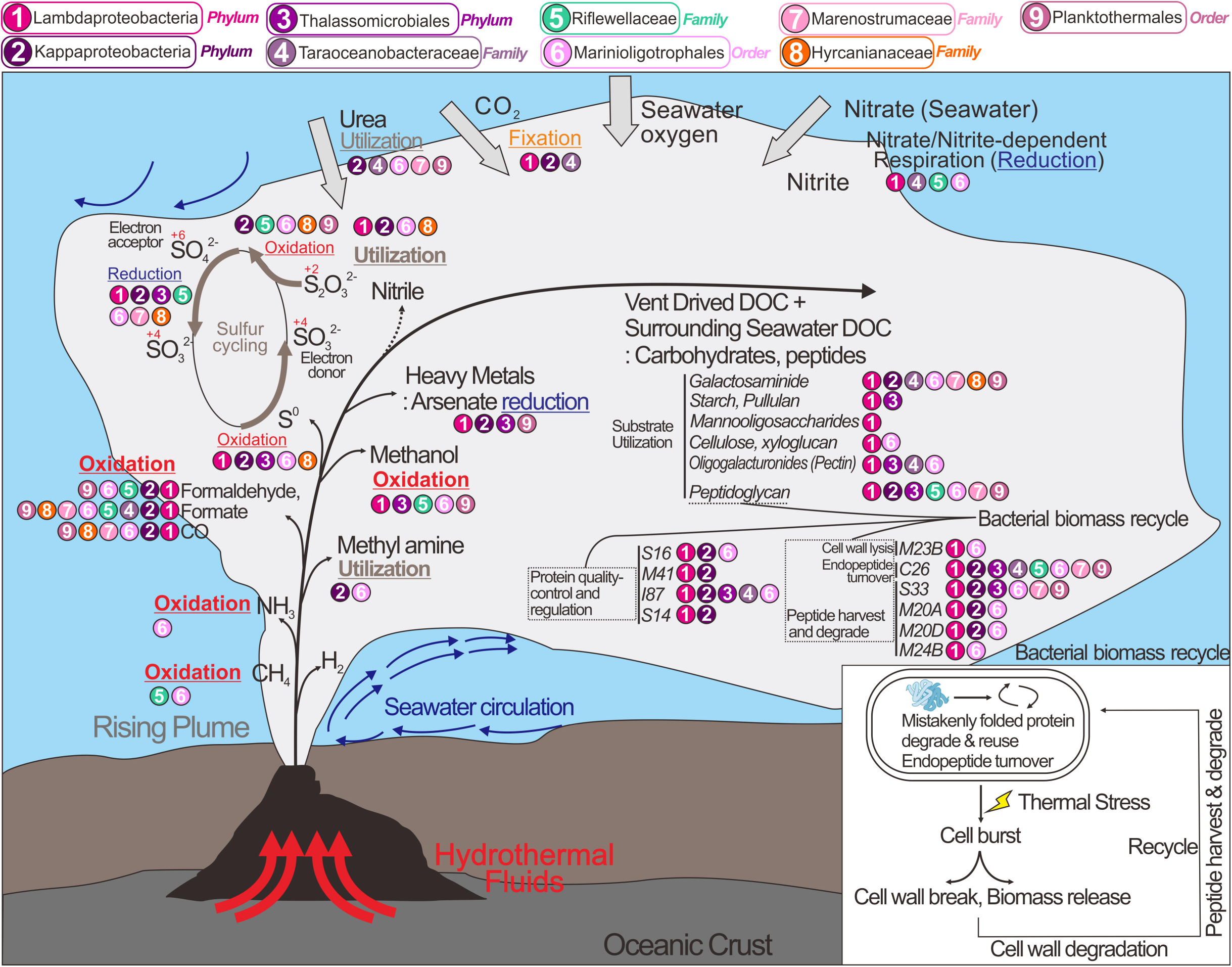
Conceptual representation of novel Proteobacteria and their ecological roles during the development and dispersal of hydrothermal plumes and the deep-sea. Ecological roles described here drive genome diversification. Hydrothermal plumes are shown to be distinct from the deep ocean water column. This distinction arises from temperature and density gradients of fluid masses. Only traits associated with biogeochemical cycling and energy metabolism are shown.

Finally, many of the novel Proteobacteria genomes encode highly-expressed genes for the breakdown of halogenated and nitrile compounds (Figs. 4, 5). To the best of our knowledge, the presence and activity of nitrile hydratases in the hydrothermal vent ecosystems has never been reported before. Although there is no direct evidence of nitrile formation in hydrothermal vents, abiotic synthesis under laboratory-based hydrothermal conditions suggests the possibility of naturally-formed nitrile-compounds^41^.

### Nitrogen and sulfur metabolism

Kappaproteobacteria, Taraoceanobacteraceae, Marinioligotrophales, Marenostrumaceae, and Planktothermales genomes encode for ureases that are highly expressed suggesting that urea may be a common source of nitrogen. Nitrogen cycling activities by Proteobacteria involve both the oxidative and reductive cycle of nitrogen. Taraoceanobacteraceae, Riflewellaceae, and Marinioligotrophales encode genes associated with ammonia oxidation (*amoABC*) and nitrite/nitrate reduction (Figs. 4, 5). No single organism possessed the entire complement of genes for denitrification, significantly no genes were observed for nitric and nitrous oxide reduction in any genomes. Only genes for the membrane-bound Nar proteins for nitrate reduction were observed, no periplasmic Nap genes for nitrate reduction were observed in any genomes. Genes for nitrite reduction included the copper-containing *nirK* (for reduction to nitric oxide) and *nirBD* and *nrfA* for dissimilatory reduction of nitrite to ammonia. Many of the Proteobacteria were associated with sulfur cycling comprising the oxidative cycle of sulfur. All proteobacterial genomes lacked genes for the oxidation of hydrogen sulfide to sulfur. However, many of them encode genes for sulfur dioxygenases (*sdo*) and reverse-dissimilatory sulfite reductases (*rdsr*) for sulfur oxidation. In particular, Kappaproteobacteria genomes encode highly active genes for complete oxidation of sulfur to sulfate including *dsrAB* for the oxidation of elemental sulfur to sulfite and *aprAB* and *sat* for the oxidation of sulfite to sulfate (Supplementary Fig. S4, S5, and Supplementary Table S10). Kappaproteobacteria, Riflewellaceae, Hyrcanianaceae, and Planktothermales genomes possessed the Sox enzyme complex for the utilization of thiosulfate. We investigated the presence of the soxCD genes in all genomes for complete oxidation of thiosulfate in lieu of disproportionation. Amongst these, Kappaproteobacteria genomes lacked soxCD suggesting that they can only disproportionate thiosulfate to elemental sulfur and sulfate while Riflewellaceae, Hyrcanianaceae, and Planktothermales can undertake complete oxidation of thiosulfate to sulfate. Kappaproteobacteria and Hyrcanianaceae genomes exhibited high levels of *sox* gene expression suggesting active utilization of thiosulfate in hydrothermal plumes (Figs 4, 5 and Supplementary Table S10). Overall, these results suggest that these novel Proteobacteria actively oxidize and cycle various nitrogen and sulfur species as nutrient and energy sources^10^.

### Metabolism of iron and other metals

Genes encoding for mineral transport enzymes for Fe (III) could be found in the Lambdaproteobacteria, Riflewellaceae, Marinioligotrophales, Marenostrumaceae, Hyrcanianaceae and Planktothermales genomes, suggesting that these Proteobacteria likely participate in the acquisition of Fe from precipitating minerals in hydrothermal plumes. Microorganisms store iron within the cell by reducing Fe (III) to Fe (II), and incorporate iron into a variety of organic compounds by forming C-Fe or S-Fe bonds, such as in metalloproteins, ferredoxins, and NADH dehydrogenase which are of significant importance to cellular activities^42^. In doing so, plume proteobacteria may serve as a part of the microbial Fe pump to scavenge and store mineral-bound Fe in biomass, to sequester Fe in the organic carbon pool after cell death, and to transfer Fe widely as plumes disperse across the oceans^43^.

Nearly all Proteobacteria contained genes for arsenate reduction (*arsC*), and many *arsC* genes have considerably high expression in hydrothermal plumes, such as in Planktothermales at Cayman, and Taraoceanobacteraceae and Kappaproteobacteria at Guaymas. Arsenic resistance and detoxification are mediated by ArsC which have previously been reported from hydrothermal environments and were found to be abundantly distributed in an iron-rich hydrothermal microbial mat from Lō’ihi Seamount, Hawai’i^44^. Since arsenic and arsenic minerals are discharged in hot, mineralized hydrothermal fluids^45^, all life forms in close proximity to the rising plume have to be resistant to elevated arsenic concentrations. To better adapt to their local environment, these hydrothermal plume proteobacterial groups have evolved to obtain both arsenic resistance and detoxification systems.

### Linking genome diversification and adaptation of functional traits

Microbial traits often evolve in close coordination with their environment. Therefore, we investigated genome diversification of individual clades of novel Proteobacteria in the context of metabolic functions that are unique to them and the distribution of gene orthologs. Amongst all novel proteobacterial lineages, Thalassomicrobiales, Kappaproteobacteria, Muproteobacteria, and Lambdaproteobacteria exhibited the strongest evidence of genome diversification and its association with environmental adaptation.

The concatenated ribosomal protein-based phylogenetic tree for Thalassomicrobiales indicated the presence of three distinct clades, namely the photic zone clade, the non-photic zone clade, and the marine subsurface clade (Fig. 8a), which are associated with the environments they inhabit. All three clades possess a unique distribution of orthologous genes with specific functions that point to their genome diversification (Fig. 8a). The photic zone clade exclusively possesses proteorhodopsin as the light-driven proton pump for energy conservation which is important for their limited chemoorganotrophic lifestyle^46^. In spite of the importance of proteorhodopsin in these organisms, they lack the ability to synthesize retinol as a chromophore^47, 48^; in line with previous reports, we hypothesize that short-chain dehydrogenases are responsible for converting other substrates (e.g., retinal or β-carotene) to retinol^47^. The aquatic β-propeller phytase (BPP) enzymes are found in marine bacteria for the mineralization of phytate to recycle phosphorus^49^. We observed the presence of BPP, phosphate-starvation-inducible protein (PsiE), alkaline phosphatase (Pho)^50^ and phospholipase only in the photic zone clade suggesting that they are more subjected to a phosphorus-limited environment and have evolved to gain a set of response mechanisms against P-starvation. Finally, the exclusive distribution of DNA photolyases (PhrB/-like) in the photic zone also indicates that Thalassomicrobiales in this environment have evolved to repair DNA damage caused by exposure to ultraviolet light^51^ (Fig. 8a).

**Figure 8.**
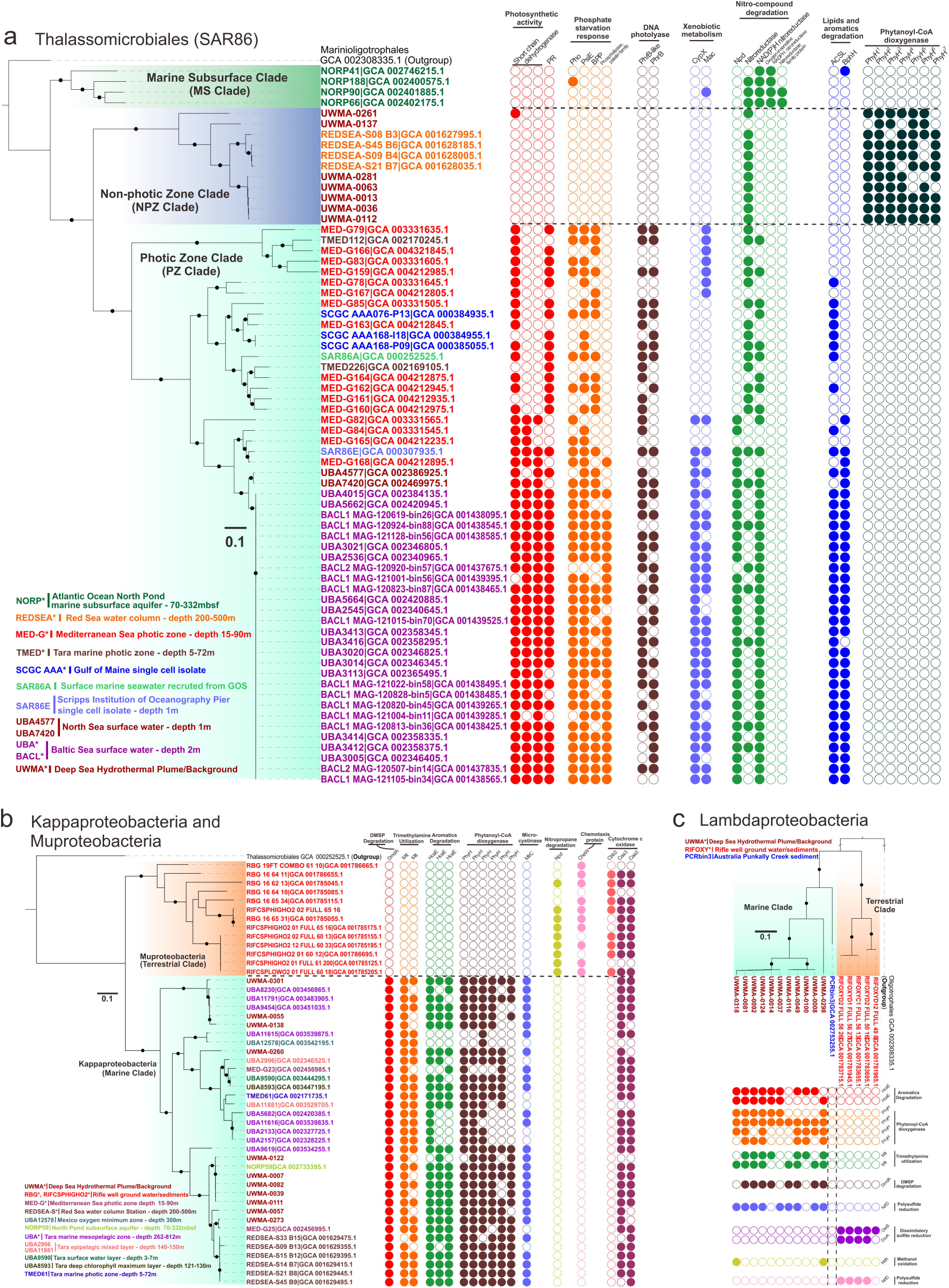
Comparison of lineage-specific protein families associated with environmental adaption in Thalassomicrobiales, Kappaproteobacteria, Muproteobacteria, and Lambdaproteobacteria populations. Phylogenetic trees were reconstructed based on concatenated 16 RPs, and branches with bootstrap values [ultrafast bootstrap (UFBoot) support values] over 90% by IQ-TREE were labeled with black dots. Each circle on the panel right to the tree indicates an ortholog group (OG) with a specific function. Filled circles represent the presence and unfilled circles represent the absence of specific traits.

Genes unique to the non-photic zone clade of Thalassomicrobiales include phytanoyl-CoA dioxygenases (PhyH) associated with the breakdown of chlorophyll. The encoding genes of PhyH on all genomes in this clade were found to be encoded in multiple copies ranging from four to seven. PhyH could hydroxylate the methyl-branched chain of phytanoyl-CoA and is required in the alpha-oxidation pathway of fatty acid metabolism to move the methyl-group from beta position to the alpha position^52^. Together with the metabolism of alcohol and aldehyde dehydrogenases and acyl-CoA synthetase, it is feasible for bacteria to transform phytol (a long-chain alcohol constituent of chlorophyll) to phytanoyl-CoA and pass the products downstream towards beta-oxidation and energy utilization^52, 53^. The exclusive distribution of PhyH in the non-photic zone clade implicates that this specific group could potentially scavenge the degradation products of chlorophyll for carbon and energy demand, which could possibly be of phytoplankton-origin from the upper ocean layers (e.g., deep chlorophyll maximum)^54^. Macrolide transporter (Mac) and cytochrome P450 (CypX) are potentially responsible for the degradation of xenobiotics (non-naturally produced compounds toxic to microbes, such as therapeutic drugs, antibiotics)^47^, and their encoding genes tend to distribute in the photic zone genomes (Fig. 8a). It also indicates that photic zone genomes possess nitronate monooxygenases (Npd) and nitroreductases for degrading nitronate and nitro-containing compounds (e.g., nitroaromatics and nitroheterocyclic compounds) (Fig. 8a). Furthermore, photic zone genomes possess additional capacities to degrade long-chain fatty acids and aromatics (Fig. 8a). This potentially indicates that xenobiotic compounds, nitro-compounds, lipids, and aromatics are universal in the surface oceans and surface microbes get are adapted for biological defense and substrate utilization. Proteobacteria inhabiting the marine subsurface are also capable of degrading nitro-compounds, as their genomes contain different types of nitroreductases, such as NAD(P)H dependent, oxygen-insensitive nitroreductases, and other nitroreductase family proteins (Fig. 8a).

The two phyla Kappaproteobacteria and Muproteobacteria represent monophyletic deep-branching clades that likely share a common ancestor. Our genome diversification analysis suggests these two lineages evolved to uniquely colonize specific environments, namely terrestrial (Muproteobacteria) and marine (Kappaproteobacteria). Kappaproteobacteria genomes encode important functional traits for utilizing elemental and energy sources in the ocean (Fig. 8b). Dimethyl sulfoniopropionate (DMSP) is a widely distributed marine algal osmolyte and is well known as a significant source of carbon and sulfur for bacterioplankton ^55^, while trimethylamine (TMA) is part of the oceanic organic nitrogen pool and produced by reduction of many marine osmolytes such as glycine betaine, trimethylamine oxide (TMAO), and choline^56^. Kappaproteobacteria genomes encode enzymes to utilize all these above-mentioned compounds. Similar to Thalassomicrobiales, Kappaproteobacteria can also potentially utilize aromatics and chlorophyll degradation products in the ocean. Furthermore, Kappaproteobacteria contain genes for microcystinase which could degrade and detoxify microcystin that is generated by marine cyanobacteria^57^. These results suggest that Kappaproteobacteria are well adapted to life in the marine water column.

In contrast, Muproteobacteria genomes were primarily sourced from terrestrial ground water and sediments and were absent in marine environments^33^. Microbes encoding chemotaxis proteins (CheW) are better adapted to sense and respond to the chemical gradients beneficial to their survival in porous underground environments. Nearly all Muproteobacteria genomes we examined possessed CheW. Muproteobacteria also encode genes to utilize nitropropane (a potential industrial hazardous compound) from terrestrial environments (Fig. 8c). Despite their existence in terrestrial environments with limited oxygen, Muproteobacteria possess cytochrome *c* oxidases and other complexes for aerobic respiration (Supplementary Table S11). Muproteobacteria genomes contain both the caa3 type (also referred to as A-type) and cbb3 type (also referred to as C-type) cytochrome *c* oxidases, while Kappaproteobacteria only contain the caa3 type. The cbb3 type cytochrome *c* oxidase has a higher oxygen binding affinity in comparison to caa3 and helps microorganisms to respire under low-oxygen conditions^58^. These results paint the picture of Muproteobacteria as a lineage of organisms that are highly adapted to survive and proliferate in the terrestrial subsurface.

Similar to Kappaproteobacteria and Muproteobacteria, the marine clade of Lambdaproteobacteria contains genes for degradation of aromatics, chlorophyll degradation products, DMSP, and TMA (Fig. 8). The functional trait of methanol oxidation is essential for utilizing methanol entrained in the hydrothermal plume as indicated (Fig. 8). Both the terrestrial and marine clades of Lambdaproteobacteria contain genes for the reduction of polysulfide, however, the enzymes are from different protein families, which suggests that they might operate differently in corresponding environments (Fig. 8). In contrast, the terrestrial clade of Lambdaproteobacteria exclusively contains the entire dissimilatory sulfite reductase gene operon (*dsr*)^59^, which suggests that they could participate in sulfite reduction to hydrogen sulfide.

## Discussion

Overall, our research provides the first comprehensive study into the ecological functions and metabolic capacities of globally distributed novel proteobacterial lineages that are abundant and active in deep-sea hydrothermal systems. Specifically, within deep-sea hydrothermal plume environments, our results suggest that novel proteobacteria have versatile metabolisms associated with chemolithotrophic activity and utilization of C1 compounds, sulfur and thiosulfate oxidation, and organotrophic activity dependent on fatty acids, aromatics, carbohydrates, and peptidases. All novel proteobacterial lineages are comprised of organisms that can respire oxygen or nitrate/nitrite.

Compared to dominant chemolithotrophic activities of the plume microbiome based on sulfur, hydrogen, and methane^60^, the metabolic versatility of these novel Proteobacteria suggests the use of specific organic substrates to enable adaptation and survival in hydrothermal plumes and surrounding environments. Bacterial cells in proximity to venting fluids could be subjected to high-temperature stress, which could cause protein unfolding^61^ and mistranslation^62^, and even cell burst. Novel Proteobacteria in this specific environment possess abundant and active peptidases for protein quality control and regulation and enzymes for endopeptide turnover. Finally, they also possess peptidoglycan degrading enzymes, peptidases for cell wall lysis, and peptide harvest and degrading systems to use organic matter which could be released to surrounding environments after cell burst. This suggests that these bacteria could actively recycle biomass within the cell and in ambient environments under hydrothermal conditions.

Although the novel proteobacteria discovered in this study are reconstructed from hydrothermal environments, they are distributed worldwide in various environments. Our study highlights how genome-based functional studies can identify novel microorganisms and their metabolic contributions, and decipher their activities associated with biogeochemical transformations and adaptive mechanisms in various environments. Genome diversification in Proteobacteria is associated with environmental adaptation^63^; and comparative genomics identifies functional traits that could potentially explain the niche-adapting mechanisms of marine/terrestrial and marine layer divisions. The evolutionary strategy adopted by Proteobacterial lineages suggests that they can flexibly modify their gene repertoire in response to substrate and energy conditions in the surrounding environments. The novel proteobacterial community discovered in the hydrothermal ecosystem are not made up of highly-adapted microbial lineages that are only limited in this environmental setting, but rather of universal lineages that have adopted strategies to live there. These findings thus open the door for further research into detecting and quantifying the wide ecological impacts of novel Proteobacteria. Our approach of studying the lineage-specific gene components will facilitate further investigations on links between genome diversification patterns and functional ecology in other microbial groups and environments.

## Methods and Materials

### Hydrothermal vent sample and metagenome sequencing

The hydrothermal vent plume, background samples, and Cayman vent fluid samples were acquired from the following cruises: R/V *New Horizon* to Guaymas Basin (July 2004), R/V *Atlantis* to Mid-Cayman Rise (Jan 2012 and Jun 2013) for Cayman Deep (*Piccard*) and Shallow (*Von Damm*), and R/V *Thomas G Thompson* to the Eastern Lau Spreading Center (ELSC) (May-Jul 2009). The detailed sampling methods and the geographic and oceanographic environmental setting information are provided elsewhere^9, 27, 29^. In brief, the plume, seawater, and vent fluid samples were collected either by Suspended Particulate Rosette (SUPR) filtration device mounted to the remotely operated vehicle or CTD-Rosette bottles^27^, and the filters (0.2 μm pore size) were preserved from microbial specimen collection. The metagenomic DNA was retrieved and sequenced accordingly by Illumina HiSeq 2000 platform (refer to previous publications^9, 27–29^). The downstream raw reads quality control was processed by Trim Galore implemented in metaWRAP v0.8.6^64^ using default settings.

### Metagenomic binning and genome refining

MetaSPAdes v3.12.0^65^ was used to assemble QC-passed reads with the settings as follows “--meta -k 21,33,55,77,99”. The QC-passed reads from the individual hydrothermal sites were combined and subjected to assembly. For samples from Mid-Cayman Rise, due to the high memory demand for large datasets, MEGAHIT v1.1.2 was used for the assembly with the following parameters “--k-list 21,33,55,77,99 -m 0.95”. The resulting assemblies (min scaffold length ≥ 1kb) and QC-passed reads were used for metagenomic binning by metaWRAP v0.8.6^64^ with self-implemented MaxBin2^66^, metaBAT^67^, and metaBAT2^68^ binning modules. A fourth-round of binning was conducted using the deep learning metagenomic binner MetaGen^69^. Finally, four sets of metagenomic bins (MAGs) were pooled together and subjected to bin dereplication, aggregation and scoring by DAS Tool with the setting as “--score_threshold 0.4”^70^, resulting in refined bins with the best quality and completeness scores.

All resulting bins representing draft quality genomes with genome completeness > 50% and contamination < 10% were further subjected to bin refinement to screen heterogeneous scaffolds potentially originated from contamination and erroneous 16S rRNA scaffolds that do not fit the genome taxonomy using RefineM v0.0.24^31^ with “gtdb_r86_protein_db” and “gtdb_r80_ssu_db”, respectively. A final round of genome refinement was conducted manually using VizBin^71^ by visualizing scaffolds and clustering them with coverage and 5-nucleotide kmer information.

### Distribution map

From each novel proteobacterial group, the longest 16S rRNA gene sequences were picked and used for comparison using BLAST against metagenomes (E-value < 1e-5) in IMG DOE metagenome database (16S rRNA Public Assembled Metagenomes accessed 2019-04-22). The BLAST hits with sequence similarity within the taxonomic division requirement were retained^72^ (only the first 500 hits were retained if there are more than 500 available hits). The IMG metagenome geographic and environmental details were parsed out and used to make the plots accordingly (*R* packages: “ggplot2”, “ggmap”, “maps” and “mapdata”).

### Phylogenetic reconstruction and genome property

The 16 ribosomal proteins (RPs, L14, L15, L16, L18, L22, L24, L2, L3, L4, L5, L6, S10, S17, S19, S3 and S8)^33^ were parsed out from each genome and aligned with our homemade archaeal and bacterial sequence databases by HMMER v3.2.1^73^ and MAFFT v7.271^74^. Only genomes containing over 4 ribosomal proteins were included in the analysis. Subsequently, the concatenated RP alignments were used for phylogenetic tree reconstruction (referred to as RP16 tree) by IQ-TREE v1.6.9^65, 75^ with the following settings “-m MFP -bb 100 -s -redo -mset WAG,LG,JTT,Dayhoff -mrate E,I,G,I+G -mfreq FU -wbtl” (“LG+I+G4” was chosen as the best-fit tree reconstruction model). The taxonomic positions of proposed proteobacterial phyla, orders and families were manually inspected in the RP16 tree, and the corresponding names were proposed accordingly. The genome characteristics of all MAGs were parsed out, which includes 1) genome phylogeny (GTDB, NCBI and manually-curated) based on Genome Taxonomy Database using GTDB-Tk v0.1.3^30^ (https://github.com/Ecogenomics/GtdbTk), 2) genome coverage, completeness and contamination, and strain heterogeneity (by CheckM^76^), 16S rRNA phylogeny and genome characteristics (The 16S phylogeny was manually checked, and those ones with incongruent taxonomy with RP16 phylogeny were filtered), and tRNA statistics (by tRNAscan-SE v2.0^77^). The 16S rRNA gene phylogenetic tree was reconstructed by aligning sequences with MAFFT v7.271^74^ followed by the construction of a phylogenetic tree building using IQ-TREE v1.6.9^65, 75^ with the settings of “-st DNA -m GTR+G4+F -bb 1000 -alrt 1000”.

### Metabolic gene annotation

We applied the biogeochemical functional trait profiler METABOLIC on reconstructed genomes^78^. Hmmsearch (HMMER 3.2.1^73^) was used to scan for potential metabolic genes from MAGs against a custom HMM database using manually curated noise cutoffs (NC). For the sulfur cycling genes, we applied manually curated trusted cutoffs (-TC option) for scanning HMMs. For *dsr* gene operon results, all results were we manually checked manually to avoid potential omissions. The custom HMM database was modified from our previous publication^33^. For detailed information, please refer to the software package METABOLIC^78^.

KEGG Orthology (KO) annotation was conducted by assigning a KO identifier to a protein using GhostKOALA^79^, KAAS^80^, and EggNOG emapper v4.5.1^81^. The KO id was assigned to a protein in the following the order 1) GhostKOALA KO, 2) KAAS KO, 3) EggNOG emapper KO and 4) EggNOG emapper COG transferred KO. As a fifth approach, we also used the NCBI-nr database (Jun 2018 released) as the reference implemented with DIAMOND v0.9.24 with default settings to annotate proteins, and the proteins were annotated with the gene product name and taxonomy of the top BLASTP hit from top 10 returned hits. If all five annotation methods produced no valid annotations, we assigned “N/A” to this protein.

### Genome functional profile and genomic comparative analysis

Reference genomes for inference of novel proteobacterial groups were downloaded by 1) searching the GTDB database (release86), 2) parsing the NCBI Assembly accession identifiers to download corresponding genomes from NCBI and 3) parsing the downloaded Genbank files for corresponding protein, genome, gene, rRNA and tRNA files. The phylogeny of reference genomes was checked by building RP16 tree using the method mentioned above.

For general functional profile analysis, all genomes were dereplicated using dRep^82^ and the genomes with over 80% genome completeness, less than 10% genome redundancy (CheckM genome characteristics; except for some lineages with few genomes, e.g., Thalassomicrobiales and Hyrcanianaceae, which also include several genomes over 70% completeness) were used. The software METABOLIC was used to obtain metabolic functional trait profiles of both reference genomes and genomes from this study^78^. The assignment of a KEGG module (a collection of KOs that could be treated as a functional unit, including pathway modules, structural complexes, functional sets and signature modules) to a proteobacterial group was conducted by firstly assigning the existence of a KEGG module compositional KO (the cutoff value for its existence in group members is 50%), and subsequently, assigning the existence of a KEGG module by the presence of all the compositional KOs (cutoff value is 75%). In addition to the KEGG KofamKOALA database^83^, a custom HMM database was also constructed and used to complement the overall functional profile (A cutoff value of 50% was used to determine the existence of a functional trait among group members).

For comparative analysis of genomes, all the available reference genomes from both GTDB and NCBI databases were downloaded and parsed using the method as mentioned above. Genomes with incorrect taxonomic assignments were removed from downstream analysis. To compare gene component differences between clades, we pulled all genome proteins together and clustered all proteins into ortholog groups (OGs) using OrthoFinder v2.2.7^84^. The OG function was annotated using EggNOG emapper and NCBI-nr database by the majority hit (over 50%) of proteins within. The comparative genomic analysis was conducted by sorting the distribution of OG among different clades using homemade Perl scripts.

### Carbohydrate-active enzyme and peptidase annotation

Proteins were parsed by hmmscan^73^ against the dbCAN2 database^85^ (dbCAN-HMMdb-V7) for annotating carbohydrate-active enzymes (CAZymes). Only the Glycoside Hydrolase (GH) and Polysaccharide Lyase (PL) annotations were retained since the function and substrate of these enzymes are better assigned. Peptidase (also including peptidase inhibitor) annotation was conducted by using DIAMOND BLASTP to search against the MEROPS database (pepunit)^86^ with the following settings: “-k 1 -e 1e-10 --subject-cover 80 --id 50”.

### Metagenomic and metatranscriptomic analysis

QC-passed metagenomic reads were used for mapping against resulting MAGs using Bowtie 2 v2.3.4.1^87^ using default settings. The normalized genome coverage was calculated by using the average coverage for all scaffolds and normalizing it with 100M reads per metagenomic dataset. QC-passed and rRNA-filtered^88^ metatranscriptomic reads were mapped against the genes of reconstructed genomes. The metric Reads Per Kilobase per Million mapped reads (RPKM) was calculated by normalizing the sequencing depth (per million reads) and the length of the gene (in kilobases). To compare the expression level of individual genes in genomes from difference environments, we also normalized RPKM values by dividing them with corresponding genome coverage. The gene product information was determined from the total annotation results accordingly.

## Data availability

Raw metagenome and metatranscriptome sequence reads are deposited in NCBI BioProject database with the accession numbers of PRJNA314399, PRJNA283159, PRJNA234377, PRJNA72707 and PRJNA283173; for detailed accession numbers refer to Supplementary Table S1. NCBI Genbank accession numbers for individual genomes could be found under the BioProject ID PRJNA522654. Additional detailed annotation results for individual genomes are available from the corresponding author on request.

## Acknowledgments

We thank the University of Wisconsin - Office of the Vice Chancellor for Research and Graduate Education, University of Wisconsin – Department of Bacteriology, and University of Wisconsin – College of Agriculture and Life Sciences for their support.

## Author contributions

Z.Z. and K.A. designed the research, conducted the analyses, and drafted the manuscript; P.Q.T. and K.K. contributed to bioinformatic analyses and data visualization. All authors reviewed the results and approved the manuscript.

## References

1. Hauptmann AL, et al. Bacterial diversity in snow on North Pole ice floes. Extremophiles 18, 945–951 (2014).

2. Zehr JP, Carpenter EJ, Villareal TA. New perspectives on nitrogen-fixing microorganisms in tropical and subtropical oceans. Trends Microbiol 8, 68–73 (2000).

3. Delmont TO, et al. Nitrogen-fixing populations of Planctomycetes and Proteobacteria are abundant in surface ocean metagenomes. Nat Microbiol 3, 804–813 (2018).

4. Huber JA, et al. Microbial population structures in the deep marine biosphere. Science 318, 97–100 (2007).

5. González JM, et al. Bacterial community structure associated with a dimethylsulfoniopropionate-producing North Atlantic algal bloom. Appl Environ Microbiol 66, 4237–4246 (2000).

6. Suzuki MT, Béjà O. An elusive marine photosynthetic bacterium is finally unveiled. Proc Natl Acad Sci U S A 104, 2561–2562 (2007).

7. Swan BK, et al. Potential for chemolithoautotrophy among ubiquitous bacteria lineages in the dark ocean. Science 333, 1296–1300 (2011).

8. Arístegui J, Gasol JM, Duarte CM, Herndld GJ. Microbial oceanography of the dark ocean’s pelagic realm. Limnol Oceanogr 54, 1501–1529 (2009).

9. Li M, Baker BJ, Anantharaman K, Jain S, Breier JA, Dick GJ. Genomic and transcriptomic evidence for scavenging of diverse organic compounds by widespread deep-sea archaea. Nat Commun 6, 8933 (2015).

10. Dick GJ. The microbiomes of deep-sea hydrothermal vents: distributed globally, shaped locally. Nat Rev Microbiol, (2019).

11. Reysenbach A-L, Banta AB, Boone DR, Cary SC, Luther GW. Biogeochemistry: microbial essentials at hydrothermal vents. Nature 404, 835 (2000).

12. Jannasch HW, Mottl MJ. Geomicrobiology of deep-sea hydrothermal vents. Science 229, 717–725 (1985).

13. Cleaves HJ. The prebiotic geochemistry of formaldehyde. Precambrian Res 164, 111–118 (2008).

14. Orita I, Yurimoto H, Hirai R, Kawarabayasi Y, Sakai Y, Kato N. The archaeon *Pyrococcus horikoshii* possesses a bifunctional enzyme for formaldehyde fixation via the ribulose monophosphate pathway. J Bacteriol 187, 3636–3642 (2005).

15. Sokolova TG, Henstra A-M, Sipma J, Parshina SN, Stams AJ, Lebedinsky AV. Diversity and ecophysiological features of thermophilic carboxydotrophic anaerobes. FEMS Microbiol Ecol 68, 131–141 (2009).

16. Lang SQ, Butterfield DA, Schulte M, Kelley DS, Lilley MD. Elevated concentrations of formate, acetate and dissolved organic carbon found at the Lost City hydrothermal field. Geochim Cosmochim Acta 74, 941–952 (2010).

17. Martin W, Russell MJ. On the origin of biochemistry at an alkaline hydrothermal vent. Philos Trans R Soc Lond B Biol Sci 362, 1887 (2007).

18. Haberstroh P, Karl D. Dissolved free amino acids in hydrothermal vent habitats of the Guaymas Basin. Geochim Cosmochim Acta 53, 2937–2945 (1989).

19. Dick G, Anantharaman K, Baker B, Li M, Reed D, Sheik C. The microbiology of deep-sea hydrothermal vent plumes: ecological and biogeographic linkages to seafloor and water column habitats. Front Microbio 4, (2013).

20. Foustoukos DI, Seyfried WE, Jr. Hydrocarbons in hydrothermal vent fluids: the role of chromium-bearing catalysts. Science 304, 1002–1005 (2004).

21. Sievert SM, Vetriani C. Chemoautotrophy at deep-sea vents: past, present, and future. Oceanography 25, 218–233 (2012).

22. Li M, Jain S, Baker BJ, Taylor C, Dick GJ. Novel hydrocarbon monooxygenase genes in the metatranscriptome of a natural deep-sea hydrocarbon plume. Environ Microbiol 16, 60–71 (2014).

23. Flores GE, et al. Inter-field variability in the microbial communities of hydrothermal vent deposits from a back-arc basin. Geobiology 10, 333–346 (2012).

24. Li Y, Liles MR, Halanych KM. Endosymbiont genomes yield clues of tubeworm success. ISME J 12, 2785–2795 (2018).

25. Sanders JG, Beinart RA, Stewart FJ, Delong EF, Girguis PR. Metatranscriptomics reveal differences in *in situ* energy and nitrogen metabolism among hydrothermal vent snail symbionts. ISME J 7, 1556 (2013).

26. Duperron S, Sibuet M, MacGregor BJ, Kuypers MM, Fisher CR, Dubilier N. Diversity, relative abundance and metabolic potential of bacterial endosymbionts in three *Bathymodiolus* mussel species from cold seeps in the Gulf of Mexico. Environ Microbiol 9, 1423–1438 (2007).

27. Lesniewski RA, Jain S, Anantharaman K, Schloss PD, Dick GJ. The metatranscriptome of a deep-sea hydrothermal plume is dominated by water column methanotrophs and lithotrophs. ISME J 6, 2257 (2012).

28. Anderson RE, et al. Genomic variation in microbial populations inhabiting the marine subseafloor at deep-sea hydrothermal vents. Nat Commun 8, 1114 (2017).

29. Sheik CS, Anantharaman K, Breier JA, Sylvan JB, Edwards KJ, Dick GJ. Spatially resolved sampling reveals dynamic microbial communities in rising hydrothermal plumes across a back-arc basin. ISME J 9, 1434 (2014).

30. Parks DH, et al. A standardized bacterial taxonomy based on genome phylogeny substantially revises the tree of life. Nat Biotechnol 36, 996–1004 (2018).

31. Parks DH, et al. Recovery of nearly 8,000 metagenome-assembled genomes substantially expands the tree of life. Nat Microbiol 2, 1533–1542 (2017).

32. Guidi L, et al. Plankton networks driving carbon export in the oligotrophic ocean. Nature 532, 465–470 (2016).

33. Anantharaman K, et al. Thousands of microbial genomes shed light on interconnected biogeochemical processes in an aquifer system. Nat Commun 7, 13219 (2016).

34. Cho JC, Giovannoni SJ. Cultivation and growth characteristics of a diverse group of oligotrophic marine Gammaproteobacteria. Appl Environ Microbiol 70, 432–440 (2004).

35. Britschgi TB, Giovannoni SJ. Phylogenetic analysis of a natural marine bacterioplankton population by rRNA gene cloning and sequencing. Appl Environ Microbiol 57, 1707–1713 (1991).

36. (!!! INVALID CITATION !!!).

37. Chen IA, et al. IMG/M: integrated genome and metagenome comparative data analysis system. Nucleic Acids Res 45, D507–D516 (2017).

38. Lombard V, Golaconda Ramulu H, Drula E, Coutinho PM, Henrissat B. The carbohydrate-active enzymes database (CAZy) in 2013. Nucleic Acids Res 42, D490–D495 (2013).

39. Forward JA, Behrendt MC, Wyborn NR, Cross R, Kelly DJ. TRAP transporters: a new family of periplasmic solute transport systems encoded by the dctPQM genes of *Rhodobacter capsulatus* and by homologs in diverse gram-negative bacteria. J Bacteriol 179, 5482–5493 (1997).

40. Fischer B, Rummel G, Aldridge P, Jenal U. The FtsH protease is involved in development, stress response and heat shock control in *Caulobacter crescentus*. Mol Microbiol 44, 461–478 (2002).

41. Rushdi AI, Simoneit BR. Condensation reactions and formation of amides, esters, and nitriles under hydrothermal conditions. Astrobiology 4, 211–224 (2004).

42. Crichton R. Iron metabolism: from molecular mechanisms to clinical consequences. John Wiley & Sons (2016).

43. Li M, Toner BM, Baker BJ, Breier JA, Sheik CS, Dick GJ. Microbial iron uptake as a mechanism for dispersing iron from deep-sea hydrothermal vents. Nat Commun 5, 3192 (2014).

44. Jesser KJ, Fullerton H, Hager KW, Moyer CL. Quantitative PCR analysis of functional genes in iron-rich microbial mats at an active hydrothermal vent system (Lo’ihi Seamount, Hawai’i). Appl Environ Microbiol 81, 2976–2984 (2015).

45. Pichler T, Veizer J, Hall GE. Natural input of arsenic into a coral-reef ecosystem by hydrothermal fluids and its removal by Fe (III) oxyhydroxides. Environ Sci Technol 33, 1373–1378 (1999).

46. Madigan MT, John M. Martinko, Kelly S. Bender, Daniel H. Buckley, and David Allan Stahl. Brock Biology of Microorganisms, Fourteenth edition edn. Pearson (2015).

47. Dupont CL, et al. Genomic insights to SAR86, an abundant and uncultivated marine bacterial lineage. ISME J 6, 1186 (2012).

48. Sabehi G, et al. New insights into metabolic properties of marine bacteria encoding proteorhodopsins. PLoS Biol 3, e273 (2005).

49. Lim BL, Yeung P, Cheng C, Hill JE. Distribution and diversity of phytate-mineralizing bacteria. ISME J 1, 321 (2007).

50. Sebastian M, Ammerman JW. The alkaline phosphatase PhoX is more widely distributed in marine bacteria than the classical PhoA. ISME J 3, 563 (2009).

51. Essen L, Klar T. Light-driven DNA repair by photolyases. Cell Mol Life Sci 63, 1266–1277 (2006).

52. Jansen GA, Wanders RJ. Alpha-oxidation. Biochimica et Biophysica Acta (BBA)-Molecular Cell Research 1763, 1403–1412 (2006).

53. Tran P, et al. Microbial life under ice: Metagenome diversity and *in situ* activity of Verrucomicrobia in seasonally ice-covered Lakes. Environ Microbiol 20, 2568–2584 (2018).

54. Weston K, Fernand L, Mills D, Delahunty R, Brown J. Primary production in the deep chlorophyll maximum of the central North Sea. J Plankton Res 27, 909–922 (2005).

55. Kiene RP, Linn LJ, Bruton JA. New and important roles for DMSP in marine microbial communities. J Sea Res 43, 209–224 (2000).

56. Lidbury IDEA, Murrell JC, Chen Y. Trimethylamine and trimethylamine N-oxide are supplementary energy sources for a marine heterotrophic bacterium: implications for marine carbon and nitrogen cycling. ISME J 9, 760 (2014).

57. Somdee T, Thunders M, Ruck J, Lys I, Allison M, Page R. Degradation of [Dha^7^] MC-LR by a Microcystin Degrading Bacterium Isolated from Lake Rotoiti, New Zealand. ISRN Microbiology 2013, (2013).

58. Bertini I, Cavallaro G, Rosato A. Cytochrome c: occurrence and functions. Chem Rev 106, 90–115 (2006).

59. Anantharaman K, et al. Expanded diversity of microbial groups that shape the dissimilatory sulfur cycle. ISME J 12, 1715–1728 (2018).

60. Anantharaman K, Breier JA, Dick GJ. Metagenomic resolution of microbial functions in deep-sea hydrothermal plumes across the Eastern Lau Spreading Center. ISME J 10, 225 (2015).

61. Day R, Bennion BJ, Ham S, Daggett V. Increasing temperature accelerates protein unfolding without changing the pathway of unfolding. J Mol Biol 322, 189–203 (2002).

62. Poole K. Stress responses as determinants of antimicrobial resistance in Gram-negative bacteria. Trends Microbiol 20, 227–234 (2012).

63. Wiedenbeck J, Cohan FM. Origins of bacterial diversity through horizontal genetic transfer and adaptation to new ecological niches. FEMS Microbiol Rev 35, 957–976 (2011).

64. Uritskiy GV, DiRuggiero J, Taylor J. MetaWRAP-a flexible pipeline for genome-resolved metagenomic data analysis. Microbiome 6, 158 (2018).

65. Nurk S, Meleshko D, Korobeynikov A, Pevzner PA. metaSPAdes: a new versatile metagenomic assembler. Genome Res 27, 824–834 (2017).

66. Wu YW, Simmons BA, Singer SW. MaxBin 2.0: an automated binning algorithm to recover genomes from multiple metagenomic datasets. Bioinformatics 32, 605–607 (2016).

67. Kang DD, Froula J, Egan R, Wang Z. MetaBAT, an efficient tool for accurately reconstructing single genomes from complex microbial communities. PeerJ 3, e1165 (2015).

68. Kang DD, et al. MetaBAT 2: an adaptive binning algorithm for robust and efficient genome reconstruction from metagenome assemblies. PeerJ 7, e7359 (2019).

69. Xing X, Liu JS, Zhong W. MetaGen: reference-free learning with multiple metagenomic samples. Genome Biol 18, 187 (2017).

70. Sieber CM, et al. Recovery of genomes from metagenomes via a dereplication, aggregation and scoring strategy. Nat Microbiol 3, 836–843 (2018).

71. Laczny CC, et al. VizBin-an application for reference-independent visualization and human-augmented binning of metagenomic data. Microbiome 3, (2015).

72. Yarza P, et al. Uniting the classification of cultured and uncultured bacteria and archaea using 16S rRNA gene sequences. Nat Rev Microbiol 12, 635–645 (2014).

73. Eddy SR. Accelerated Profile HMM Searches. PLoS Comput Biol 7, e1002195 (2011).

74. Katoh K, Standley DM. MAFFT: iterative refinement and additional methods. Multiple Sequence Alignment Methods, 131–146 (2014).

75. Nguyen L-T, Schmidt HA, von Haeseler A, Minh BQ. IQ-TREE: a fast and effective stochastic algorithm for estimating maximum-likelihood phylogenies. Mol Biol Evol 32, 268–274 (2014).

76. Parks DH, Imelfort M, Skennerton CT, Hugenholtz P, Tyson GW. CheckM: assessing the quality of microbial genomes recovered from isolates, single cells, and metagenomes. Genome Res 25, 1043–1055 (2015).

77. Lowe TM, Chan PP. tRNAscan-SE On-line: integrating search and context for analysis of transfer RNA genes. Nucleic Acids Res 44, W54–W57 (2016).

78. Zhou Z, Tran P, Liu Y, Kieft K, Anantharaman K. METABOLIC: A scalable high-throughput metabolic and biogeochemical functional trait profiler based on microbial genomes. bioRxiv, 761643 (2019).

79. Kanehisa M, Sato Y, Morishima K. BlastKOALA and GhostKOALA: KEGG tools for functional characterization of genome and metagenome sequences. J Mol Biol 428, 726–731 (2016).

80. Moriya Y, Itoh M, Okuda S, Yoshizawa AC, Kanehisa M. KAAS: an automatic genome annotation and pathway reconstruction server. Nucleic Acids Res 35, W182–W185 (2007).

81. Huerta-Cepas J, et al. eggNOG 4.5: a hierarchical orthology framework with improved functional annotations for eukaryotic, prokaryotic and viral sequences. Nucleic Acids Res 44, D286–D293 (2016).

82. Olm MR, Brown CT, Brooks B, Banfield JF. dRep: a tool for fast and accurate genomic comparisons that enables improved genome recovery from metagenomes through de-replication. ISME J 11, 2864 (2017).

83. Aramaki T, et al. KofamKOALA: KEGG ortholog assignment based on profile HMM and adaptive score threshold. bioRxiv, 602110 (2019).

84. Emms DM, Kelly S. OrthoFinder: solving fundamental biases in whole genome comparisons dramatically improves orthogroup inference accuracy. Genome Biol 16, 157 (2015).

85. Zhang H, et al. dbCAN2: a meta server for automated carbohydrate-active enzyme annotation. Nucleic Acids Res 46, W95–W101 (2018).

86. Rawlings ND, Barrett AJ, Finn R. Twenty years of the MEROPS database of proteolytic enzymes, their substrates and inhibitors. Nucleic Acids Res 44, D343–D350 (2016).

87. Langmead B, Salzberg SL. Fast gapped-read alignment with Bowtie 2. Nat Methods 9, 357 (2012).

88. Kopylova E, Noé L, Touzet H. SortMeRNA: fast and accurate filtering of ribosomal RNAs in metatranscriptomic data. Bioinformatics 28, 3211–3217 (2012).

89. Letunic I, Bork P. Interactive Tree Of Life (iTOL): an online tool for phylogenetic tree display and annotation. Bioinformatics 23, 127–128 (2007).

