## Supplementary material for "Genome diversification in globally distributed novel marine Proteobacteria is linked to environmental adaptation": Supplementary_Figure_S1.16S_novel_proteobacteria_and_All_ref_16S_over300.mdf.delete_errors.rrna.mafft.treefile.nwk.pdf.cdr.pdf

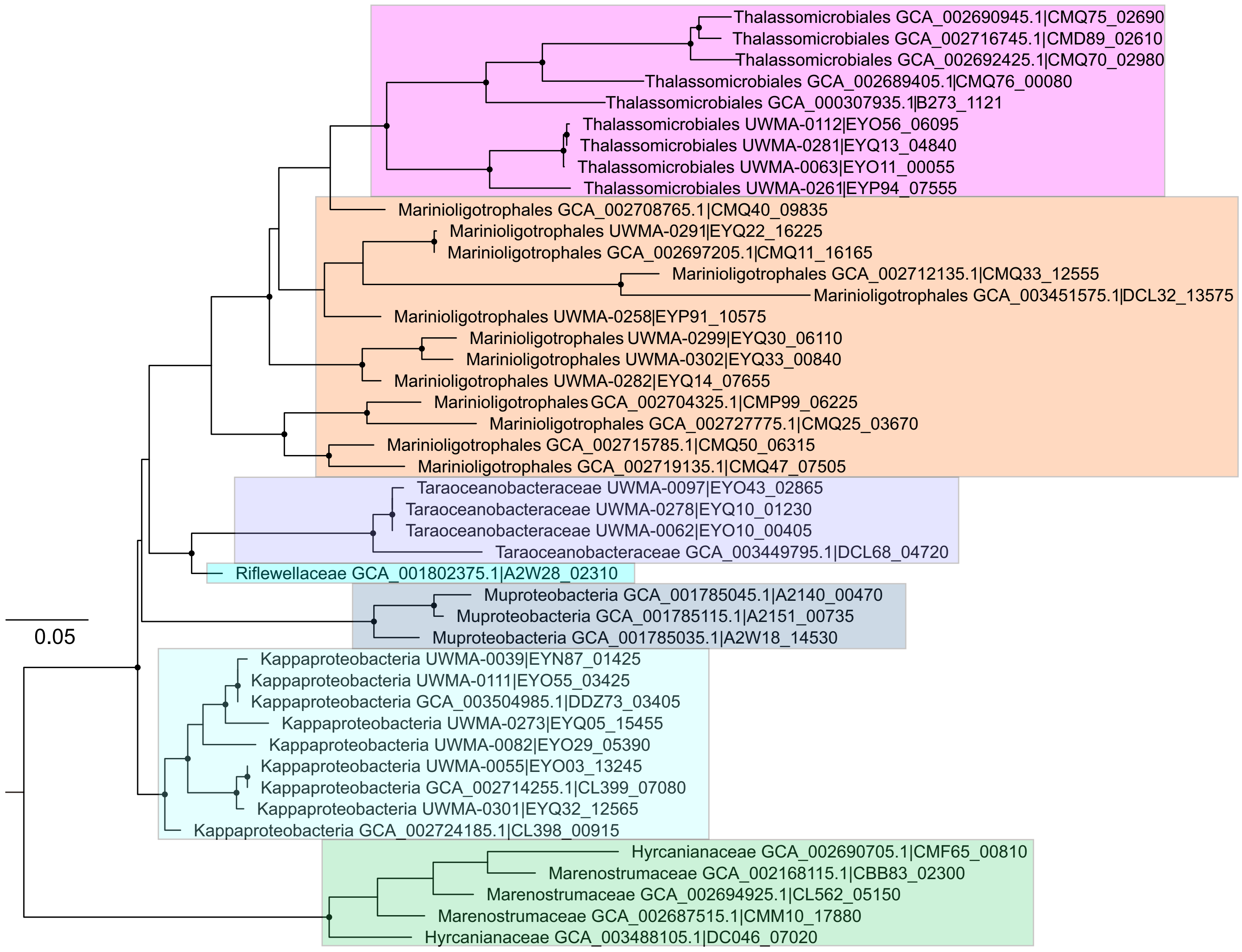


**Figure S1.** The 16S rRNA gene tree of novel Proteobacteria genomes from this study and other reference genomes. The 16S rRNA genes (over 300bp) were parsed from the genomes and aligned by MAFFT v7.271. Phylogenetic tree was constructed by IQ-TREE v1.6.9 using settings as follows: "-m GTR+G4+F -bb 1000 -alrt 1000". Potential erroneous 16S rRNA gene sequences were manually checked and screened from the tree. The black dots on the branch node indicate bootstrap values [ultrafast bootstrap (UFBoot) support values] over 90%.
