## Supplementary material for "Genome diversification in globally distributed novel marine Proteobacteria is linked to environmental adaptation": Supplementary_Figure_S2.All_proteobacteria_RP_tree.pdf

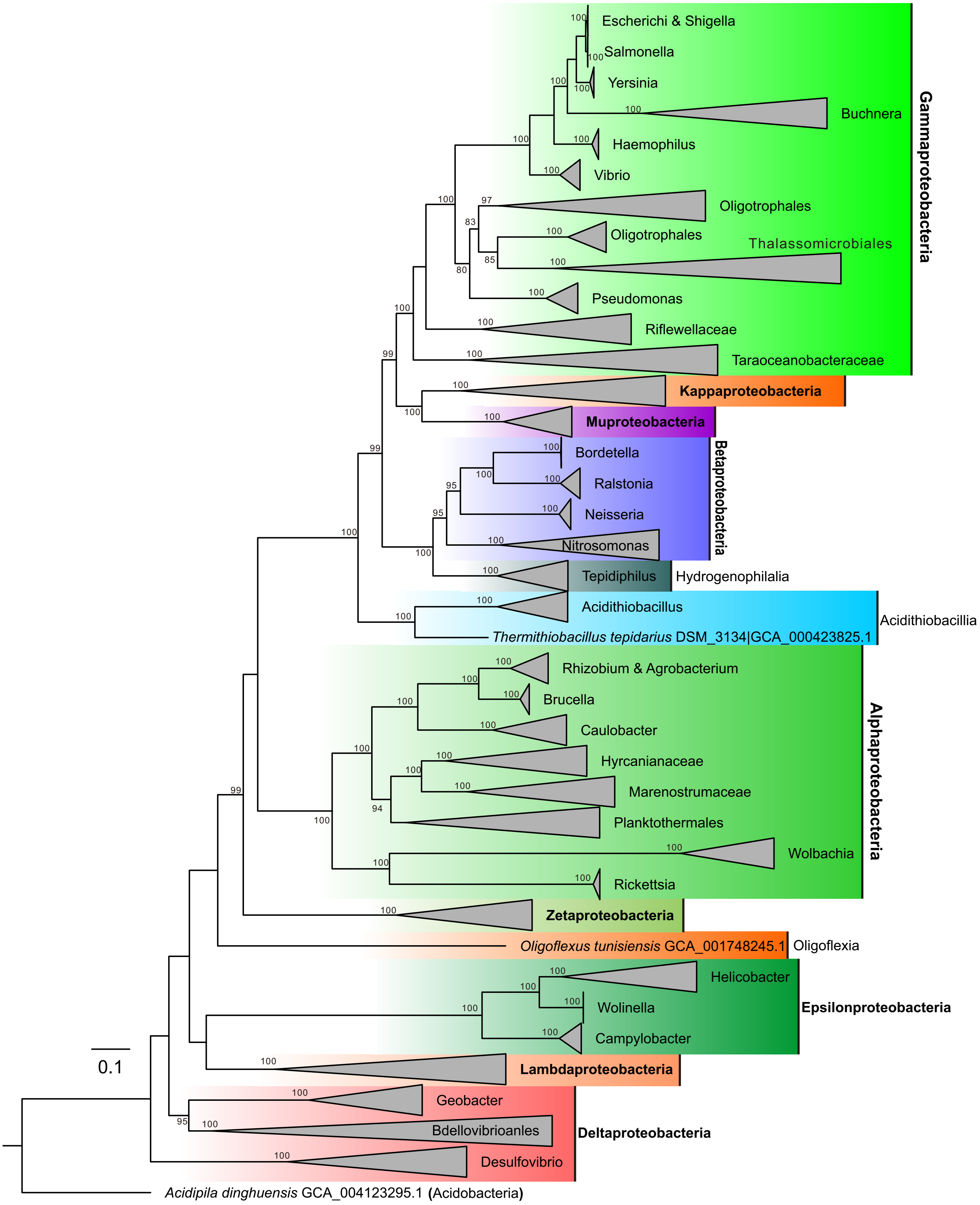


**Figure S2.** The concatenated ribosomal protein (RP) tree of novel Proteobacteria. The 16 RPs (including rpL14, rpL15, rpL16, rpL18, rpL22, rpL24, rpL2, rpL3, rpL4, rpL5, rpL6, rpS10, rpS17, rpS19, rpS3 and rpS8) were parsed from representative Proteobacteria genomes, aligned by MAFFT v7.271 individually, and then concatenated. The phylogenetic tree was reconstructed using IQ-TREE v1.6.9 with the following settings: "-mset WAG,LG,JTT,Dayhoff -mrate E,I,G,I+G -mfreq FU -wbtl". Only genomes that contain at least 4 RPs were used to build the phylogenetic tree.
