## Supplementary material for "Genome diversification in globally distributed novel marine Proteobacteria is linked to environmental adaptation": Supplementary_Figure_S3.RP_Tree_concat_over4rp.mdf.fasta.treefile.nwk.pdf.cdr.pdf

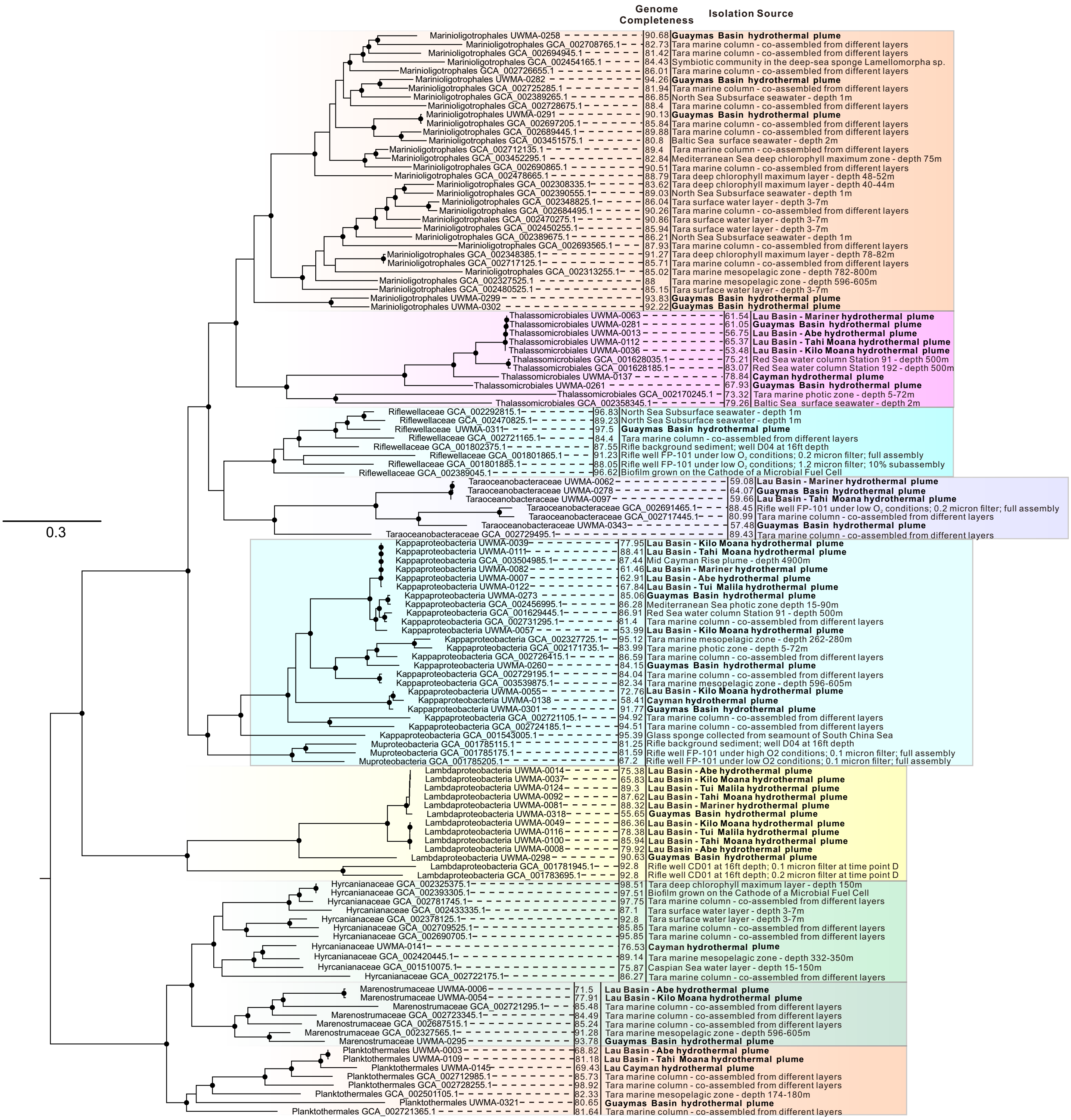


**Figure S3.** The concatenated ribosomal protein (RP) tree of novel Proteobacteria genomes from this study and other reference genomes (Genome Completeness over 80%, except for some lineages which include genomes over 70% completeness). The 16 RPs (including rpL14, rpL15, rpL16, rpL18, rpL22, rpL24, rpL2, rpL3, rpL4, rpL5, rpL6, rpS10, rpS17, rpS19, rpS3 and rpS8) were parsed from the genomes, aligned by MAFFT v7.271 individually, and then concatenated. The phylogenetic tree was reconstructed by IQ-TREE v1.6.9 using settings as follows: "-mset WAG,LG,JTT,Dayhoff -mrate E,I,G,I+G -mfreq FU -wbtl". Only the genomes that contain at least 4 RPs were used to build the phylogenetic tree. The potentially wrong assignment of taxonomy of certain genomes was screened by manual checking. The closed black circles on the branch nodes indicate bootstrap values [ultrafast bootstrap (UFBoot) support values] over 90% by IQ-TREE.
