## Supplementary material for "Genome diversification in globally distributed novel marine Proteobacteria is linked to environmental adaptation": Supplementary_Figure_S4.Gene_operon.cdr.pdf

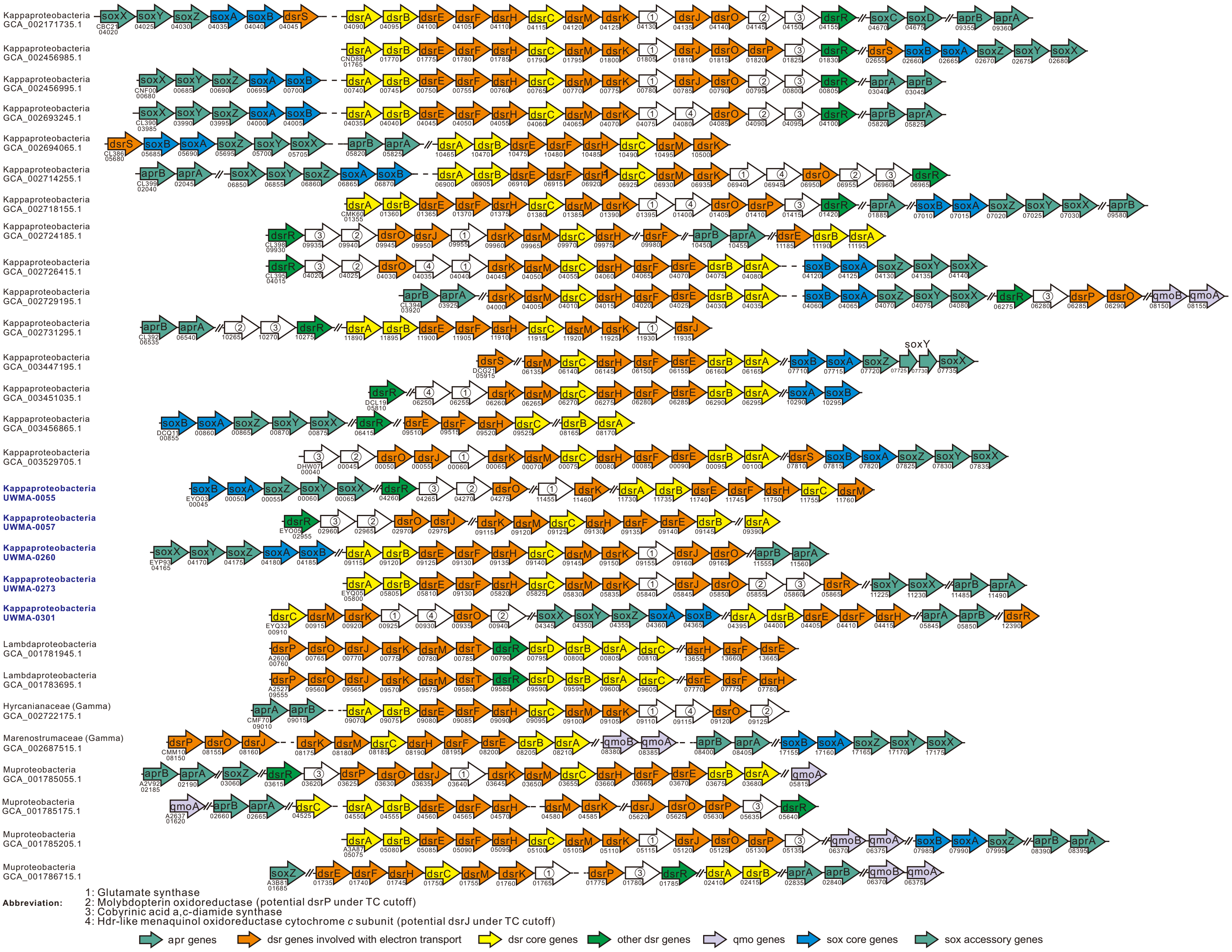


**Figure S4**. Gene arrangement of operons associated with dissimilatory sulfur metabolism. Each protein was parsed out by searches against manually curated custom HMM databases with trusted cutoffs (TC). All genes are shown on the positive strand. Direction of arrows does not indicate the real open reading frame direction along the scaffold, and the length of the arrows do not correspond to gene size. Numbers indicate gene IDs in the corresponding Genbank files on the NCBI database.
