## Supplementary material for "Genome diversification in globally distributed novel marine Proteobacteria is linked to environmental adaptation": Supplementary_Figure_S5.dsrAB.hits_and_ref.faa.mafft.gt25perc.treefile.cdr.pdf

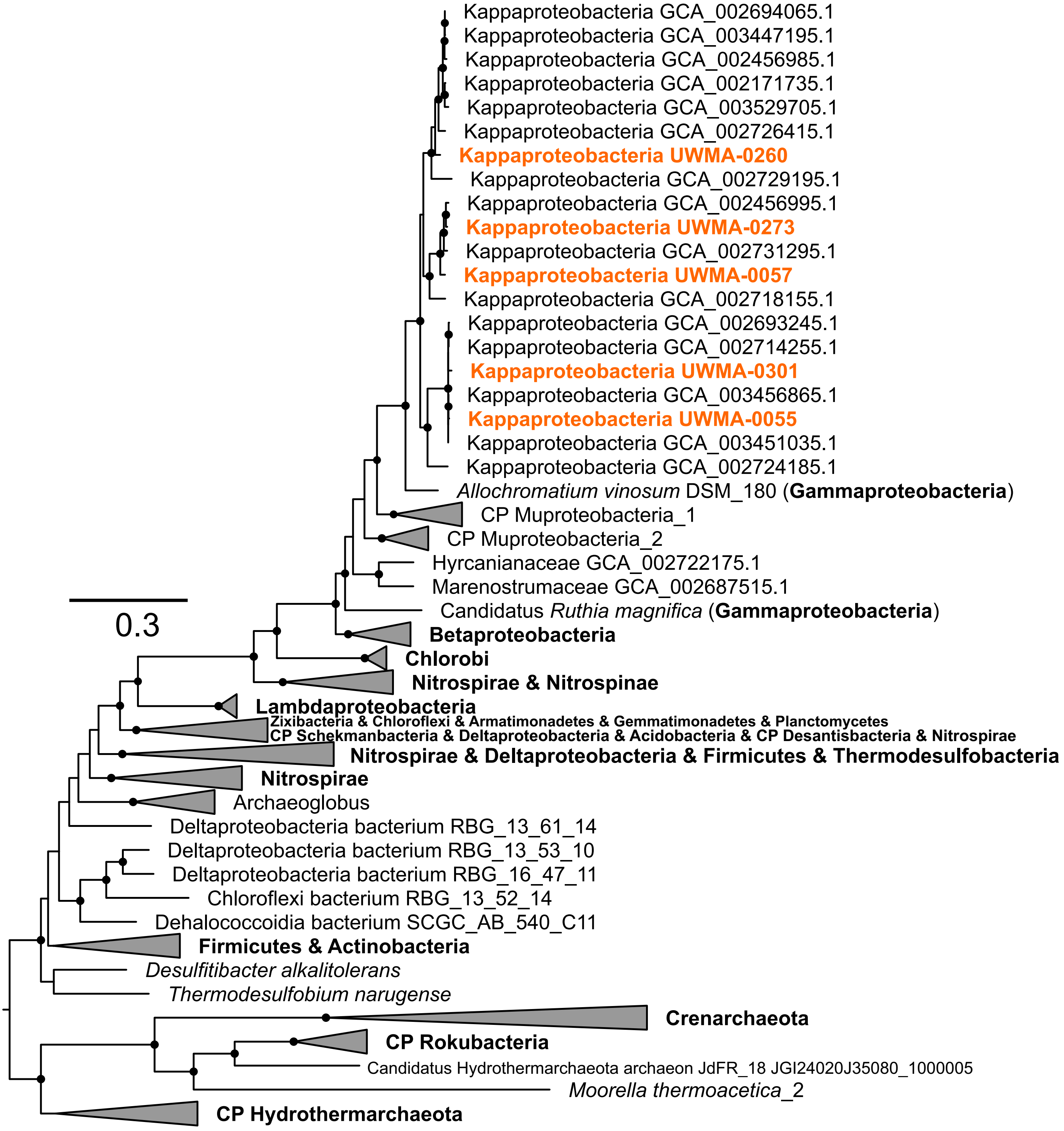


**Figure S5**. Concatenated phylogenetic tree of *dsrAB* proteins. DsrA and DsrB proteins were aligned with reference sequences independently and concatenated. The concatenated protein alignment was trimmed with gapthreshold of 25% using trimAl v1.2. The phylogenetic tree was reconstructed by IQ-TREE v1.6.9 with settings as described in the methods. Branches with over 90% UFBoot bootstrap values were labeled with closed circles. Genomes from this study were highlighted in bold.
